# A pipeline for natural small molecule inhibitors of endoplasmic reticulum stress

**DOI:** 10.1101/2022.02.20.481203

**Authors:** Daniela Correia da Silva, Patrícia Valentão, Paula B. Andrade, David M. Pereira

## Abstract

The homeostasis of eukaryotic cells is inseverable of that of the endoplasmic reticulum (ER). The main function of this organelle is the synthesis and folding of a significant portion of cellular proteins, while also being the major calcium reservoir of the cell. Upon unresolved ER stress, a set of stress response signaling pathways that are collectively labeled as the unfolded protein response (UPR) is activated. Prolonged or intense activation of this molecular machinery may be deleterious. It is known that compromised ER homeostasis, and consequent UPR activation, characterize the pathogenesis of neurodegenerative disease.

In an effort to discover new small molecules capable of countering ER stress, we subjected a panel of over 100 natural molecules to a battery of assays designed to evaluate several hallmarks of ER stress. The effect of the compounds on calcium homeostasis, key gene and protein expression, and levels of protein aggregation were evaluated in fibroblasts, and subsequently in neuronal cells. This framework resulted in the identification of several bioactive molecules capable of countering ER stress and deleterious events associated to it, among which delphinidin stands out as the most promising candidate against neurodegeneration.

## 1. Introduction

The endoplasmic reticulum (ER) plays a substantial role in the upkeep of the proteostasis of any eukaryotic cell. Upon unresolved ER stress, several stress response signaling pathways are activated in this organelle. These pathways are collectively termed as the unfolded protein response (UPR) and aim to restore reticular homeostasis, but may be deleterious upon sustained or intense activation. Shortly, the UPR relies on three major signaling branches, each initiated by its respective sensor: the protein kinase RNA-like endoplasmic reticulum kinase (PERK), inositol-requiring enzyme 1 (IRE1) or the activating transcription factor 6 (ATF6) [1, 2].

Disturbances in the secretory pathway of proteins and consequent stress upon the ER are hallmarks of several disease etiologies, including those of neurodegenerative nature. Presently, neurodegenerative diseases have a high prevalence around the world, and, as the population ages, their incidence is expected to increase [1, 3, 4]. Since pharmacological intervention by modulating ER stress has shown promising results against neurodegeneration in *in vitro* and *in vivo* studies [5–8], we consider the development of ER-based strategies against neurodegenerative diseases a necessary step to overcome the burden that these diseases represent to society.

Natural products have been used as medicinal agents from immemorial times and remain the best source of new drugs and drugs leads to modern medicine [9]. This chemical space has been gifted with immense structural diversity and has been evolutionarily tailored to provide drug-like molecules, displaying multiple chemotypes and pharmacophores [9, 10]. Furthermore, it is known that the time when pharmaceutical companies decided to reduce natural product research projects correlates with decreased numbers of new drugs entering the market [10]. A literature survey on nature-derived ER modulators [2] shows the importance of the natural product chemical space for this end, including molecules as distinct as basiliolides [11], fisetin [12–14], berberine [15–17], withaferin A [18, 19], agelasine B [20], cephalostatin 1 [21, 22], and hydroxytyrosol [23]. Notably, most of these molecules are associated to ER stress induction, while hydroxytyrosol and berberine are reported to ameliorate ER stress.

Herein we propose a pipeline for the identification of potential ER stress inhibitors among a library of natural products.

## 2. Materials and Methods

### 2.1. Chemical library and reagents

The compounds 5,7,8-trihydroxyflavone, 5-deoxykaempferol, acacetin, apigenin, apigetrin, β-escin, chrysoeriol, chrysophanol, coumarin, cyanidin, delphinidin, diosmetin, diosmin, ellagic acid, eriocitrin, eriodictyol, eriodictyol-7-*O*-glucoside, eupatorin, ferulic acid, fisetin, flavanone, galangin, gallic acid, genkwanin, gentisic acid, guaiazulene, herniarin, hesperidin, homoeriodictyol, homoorientin, homovanillic acid, hyoscyamine, isorhamnetin, isorhamnetin-3-*O*-glucoside, isorhamnetin-3-*O*-rutinoside, isorhoifolin, juglone, kaempferide, kaempferol, kaempferol-3-*O*-rutinoside, kaempferol-7-*O*-glucoside, kaempferol-7-*O*-neohesperidoside, linarin, liquiritigenin, lutein, luteolin, luteolin tetramethylether, luteolin-3’,7-di-*O*-glucoside, luteolin-4’-*O*-glucoside, luteolin-7-*O*-glucoside, malvidin, mangiferin, maritimein, myricetin, myricitrin, myrtillin, naringenin-7-*O*-glucoside, narirutin, oleuropein, orientin, pelargonidin, pelargonin, phloroglucinol, pyrogallol, quercetin-3-*O*-(−6-acetylglucoside), quercetagetin, quercetin-3,3’,4’,7-tetramethylether, quercetin-3,4’-dimethylether, quercetin-3-methylether, quercetin-3-*O*-glucuronide, rhoifolin, robinin, saponarin, scopolamine, sennoside A, sennoside B, sulfuretin, tectochrysin, tiliroside, verbascoside and vitexin were purchased from Extrasynthese (Genay, France). The molecules (-)-norepinephrine, (+/-)-dihydrokaempferol, 1,4-naphtoquinone, 18α-glycyrrhetinic acid, 3,4-dihydrobenzoic acid, 3,4-dimethoxycinnamic acid, 3-hydroxybenzoic acid, 4-hydroxybenzoic acid, 5-methoxypsoralen, ajmalicine, berberine, betanin, betulin, boldine, catechol, cholesta-3,5-diene, chrysin, cynarin, daidzein, emodin, galanthamine, genistein, guaiaverin, hesperetin, hydroquinone, lupeol, myristic acid, naringenin, naringin, *p*-coumaric acid, phloridzin, pinocembrin, plumbagin, quercetin, quercetin-3-*O*-β-D-glucoside, quercitrin, rosmarinic acid, rutin and silibinin were acquired from Sigma-Aldrich (St. Louis, MO, USA). 5,8-Dihydroxy-1,4-naphtoquinone and caffeine were from Fluka (Buchs, Switzerland), vanillin was supplied by Vaz Pereira (Santarém, Portugal), vicenin-2 was purchased from Honeywell (Charlotte, NC, USA). Chlorogenic acid was from PhytoLab (Vestenbergsgreuth, Germany). Cinnamic acid was obtained through Biopurify (Chengdug, China). Swertiamarin was from ChemFaces (Wuhan, China).

Minimum Essential Medium (MEM), Dulbecco’s Modified Eagle Medium/Nutrient Mixture F-12 (DMEM/F-12), fetal bovine serum, penicillin/streptomycin solution (penicillin 5000 units/mL and streptomycin 5000 µg/mL), trypsin-EDTA (0.25%), Qubit^®^ RNA IQ assay kit and Qubit^®^ RNA HS assay kit, the SuperScript™ IV VILO™ MasterMix, the Qubit^®^ Protein Assay Kit, the anti-BiP mouse monoclonal antibody and the anti-mouse secondary antibody were obtained from Invitrogen (Grand Island, NE, USA). Dimethyl sulfoxide (DMSO) was acquired from Fisher Chemical (Loughborough, UK). Isopropanol was obtained from Merck (Darmstadt, Germany). 3-(4,5-Dimethylthiazol-2-yl)-2,5-diphenyltetrazolium Bromide (MTT), calcium ionophore A23187, thapsigargin, RNAzol^®^, chloroform, isopropanol, diethyl pyrocarbonate (DEPC), KAPA SYBR^®^ FAST qPCR Kit Master Mix (2X) Universal, Aβ_25-35_, 2-(4-amidinophenyl)-6-indolecarbamidine dihydrochloride (DAPI), protease inhibitor cocktail, phosphatase inhibitor cocktail, acrylamide/bis-acrylamide 30% solution, bromophenol blue, glycerol, sodium dodecylsulfate (SDS), glycine, Trizma^®^ base, Trizma^®^ hydrochloride, sodium deoxycholate, sodium chloride, potassium phosphate monobasic, potassium chloride, sodium phosphate dibasic, sodium bicarbonate, D-glucose and Triton X-100 were purchased from Sigma-Aldrich (St. Louis, MO, USA), while formaldehyde was from Bio-Optica (Milan, Italy). The fluorescent probe Fura-2/AM, as well as the anti-GAPDH rabbit monoclonal antibody and the anti-rabbit secondary antibody, were obtained from Abcam (Cammbridge, UK). Thioflavin T was from Alfa Aesar (Kandel, Germany). The WesternBright^®^ ECL HRP was supplied by Advansta (Menlo Park, CA, USA). Trans-Blot Turbo Mini 0.2 µm Nitrocellulose Transfer Packs were acquired from Bio-Rad (Hercules, CA, USA).

### 2.2. Chemometric analysis

Since it is the most detailed and up to date database of annotated natural products [24], we started from the COCONUT database (https://coconut.naturalproducts.net), and created a data frame comprising all the molecules under study, where each row corresponded to a different molecule and columns to features. Among all the original features, we have decided to keep only 48, as the remaining ones did not contain relevant information for this work (SMILE notation, fragments, among others). **Table S1** contains a list of all variables kept and their description. All data was cleaned and explored using Python 3.9. For preprocessing data, we have used Simple Imputer, Standard Scaler, Ordinal Encoder and Label Encoder from Scikit-learn [25]. Dimensionality reduction was conducted using Scikit-learn [25] library (Principal Component Analysis [PCA], Multidimensional Scaling [MDS], t-distributed Stochastic Neighbor Embedding [TSNE] and Uniform Manifold Approximation and Projection (UMAP) used the UMAP library [26]. For t-SNE we have used number of neighbors of 5, 13 and 45 and minimum distance of 0.01, 0.1 and 0.5. In the case of UMAP we have iterated through perplexity of 5, 20, 50 and 100 and number of iterations of 300, 900, 1500 and 2000. Unless otherwise specified, specifications of each algorithm were set for default. Graphics in **Figures 1**, **7**, **8** and **S3** were created using Seaborn or MatPlotLib and the remaining ones in GraphPad Prism 8.

**Fig. 1.**
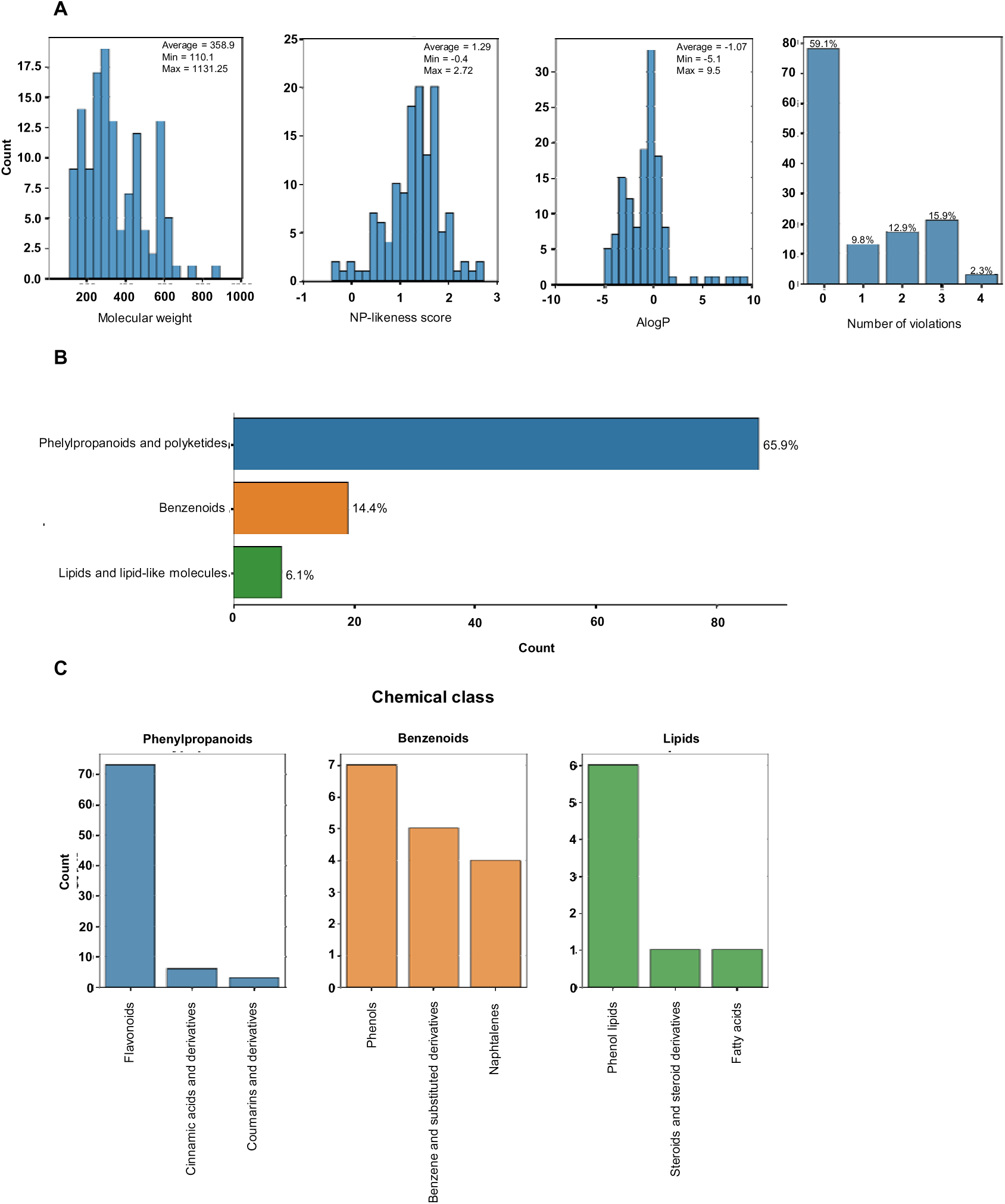
Physico-chemical properties of the library used, and distribution of the chemical (super)classes of the molecules present in the library. [2-column]

### 2.3. Cell culture conditions

MRC-5 fibroblasts (European Collection of Authenticated Cell Cultures, Porton Down Salisbury, UK) were cultured in MEM and SH-SY5Y (American Type Culture Collection, LGC Standards S.L.U., Spain) cells were cultured in DMEM/F-12. Both mediums contained 10% FBS and 1% penicillin/streptomycin and both cell lines were maintained at 37 °C with 5% CO_2_.

### 2.4. Viability assays

For viability assessment, MRC-5 fibroblasts were seeded at a density of 2 x 10^4^ cells/well. After 24 h the cells were incubated with the compounds of interest. 24 h later, the wells are aspirated, the medium was replaced by MTT at 0.5 mg/mL and incubated for 2 h. At the end of this period, the solution was discarded and the formazan crystals in the wells dissolved in 200 µL of a 3:1 DMSO:isopropanol solution. The absorbance at 560 nm was read in a Thermo Scientific™ Multiskan™ GO microplate reader.

Results are presented as the percentage of control and correspond to the mean ± SEM of, at least, three independent experiments, each of them performed in triplicate.

### 2.5. Cytosolic calcium level determination

MRC-5 fibroblasts were plated on black bottomed-96-well plates at the density of 2 x 10^4^ cells/well. After 24 h, the fluorescent probe Fura-2/AM was added at 5 µM and incubated for 1 h. Then they were incubated with the selected nontoxic compounds in HBSS. The calcium ionophore A23187 was added at a final concentration of 5 µM after 2 h. Finally, 2 h later the fluorescence was read at 340/505 and 380/505 on a Cytation™ 3 (BioTek (Vermont, MA, USA)) multifunctional microplate reader. Data analysis was performed considering the ratio F_340/505_/F_380/505_. Results are presented as fold decrease *vs* positive control and represent the mean ± SEM of, at least, three independent experiments, each of them performed in triplicate.

### 2.6. RNA extraction and quantification, cDNA synthesis and RT-qPCR reaction

Cells were seeded at 1.6 x 10^5^ cells/well in 12-well plates, left at 37 °C for 24 h and then incubated with the compounds that displayed positive results in previous assays. After 2 h, thapsigargin (Tg) was used as a positive control and added at a final concentration of 3 µM, and a period of 16 h of incubation ensued. The cells were lysed by aspirating the culture medium and adding 500 µL of PureZOL RNA isolation reagent, pipetting up and down multiple times. The lysate was kept at room temperature for 5 min and then RNA extraction ensued.

The extraction was performed by phase separation, by adding 100 µL of chloroform, shaking, incubating for 5 min and centrifuging at 12000 g for 15 min at 4 °C. The aqueous phase was collected and 250 µL of isopropyl alcohol were added, following 5 min of incubation and new centrifugation at 12000 g for 10 min, at 4°C. The supernatant was then discarded, and the pellet was washed with 75% ethanol, vortexed, centrifuged at 7500 g for 5 min at 4 °C, air dried and resuspended in DEPC-treated water. The RNA in the sample was quantified resorting to the Qubit^®^ RNA HS assay kit. Sample integrity was then evaluated with the Qubit^®^ RNA IQ assay kit, and, if in appropriate conditions, 1 µg was converted to cDNA using the SuperScript™ IV VILO™ MasterMix.

RT-qPCR reaction was performed in the following thermal cycling conditions: 3 min at 95 °C, 40 cycles of 95 °C for 3 s (denaturation), gene-specific temperature for 20 s (annealing temperatures listed on **Table 1**) and 20 s at 72 °C (extension). The employed mastermix was KAPA SYBR^®^ FAST qPCR Kit Master Mix (2X) Universal. The reaction was conducted on a qTOWER3 G (Analytik Jena AG, Germany), and the data were analyzed on the software supplied along with the equipment (qPCRsoft 4.0). Primers for target genes were designed on Primer BLAST (NCBI, Bethesda, MD, USA) and synthesized by Thermo Fisher (Waltham, MA, USA). The respective nucleotide sequences are presented on **Table 1**. Product specificity was checked with melting curves. *GAPDH* was selected as reference genes for normalization of expression. All the RT-qPCR reactions were performed in duplicate, and the experiments were repeated, at least, four times. Results are presented as mean ± SEM.

**Table 1.**
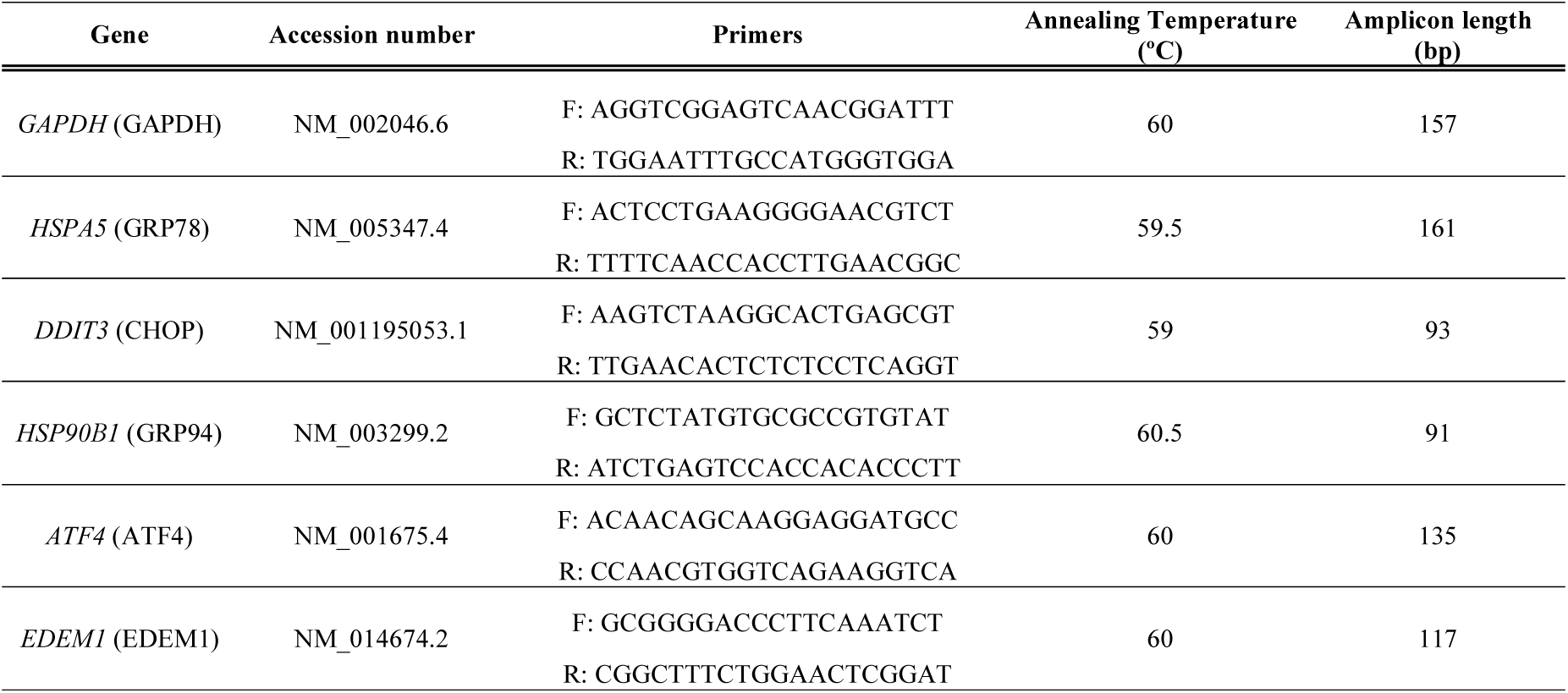
Data concerning analyzed genes and designed primer sequences.

### 2.7. Quantification and imaging of protein aggregates

On black bottomed-96-well plates, MRC-5 and SH-SY5Y cells were seeded at the densities of 2 x 10^4^ and 3 x 10^4^ cells/well, respectively. In the following day, they were incubated with the selected molecules in the presence of Aβ_25-35_ at 10 µM. After an incubation period of 24 h, cells were washed with HBSS and incubated for 30 min with thioflavin T at 5 µM (prepared in HBSS). At this point, fluorescence was read at 450/482 nm on a Cytation™ 3 (BioTek (Vermont, MA, USA)) multifunctional microplate reader. Results represent the fold decrease on fluorescence compared to the positive control. We present the mean ± SEM of, at least, three independent assays, individually performed in triplicate.

To image protein aggregates, cells were seeded on clear bottomed-96-well plates at the aforementioned density and treated with the molecules of interest for 24 h in the presence of Aβ_25-35_ at 10 µM. Cells were fixed on 4% formaldehyde at room temperature for 15 min and incubated with 10 µM thioflavin T for 60 min. The counter staining was performed with DAPI at 0.25 µg/mL, for 30 min. Finally, three washing steps with HBSS for 5 min were carried out and cells were imaged under an Eclipse Ts2R-FL (Nikon) fluorescence microscope equipped with a Retiga R1 camera. A FITC filter was employed to observe thioflavin T and DAPI staining was imaged using a DAPI filter. Images were analyzed resorting to the CellProfiler software version 4.2.1 (Broad Institute of MIT and Harvard, Cambridge, MA).

### 2.8. Western blotting analysis

MRC-5 and SH-SY5Y cells were seeded in 6-well plates at the densities of 3.2 x 10^5^ and 4.8 x 10^5^ cells/well. After 24 h, they were incubated with compounds selected as promising in previous experiments. After 2 h, Tg at 3 µM was added, and the plates were left in the incubator for another 22 h. Ended this time frame, cell samples were lysed in RIPA buffer containing 1% protease inhibitor cocktail and 1% phosphatase inhibitor cocktail for 15 min at 4 °C, and subsequently centrifuged at 14000 g for 15 min to remove cell debris. The supernatant was then collected, and the protein content was determined resorting to the Qubit^®^ Protein Assay Kit.

The following step was to submit 30 µg of each sample, previously denatured at 76 °C for 10 min, to 10% SDS-PAGE electrophoresis, in a gel containing 10 % acrylamide/bisacrylamide. The proteins were then transferred to a nitrocellulose membrane on a Trans-Blot^®^ Turbo (Bio-Rad; Hercules, CA, USA). The membrane was blocked in a PBS solution containing 5% non-fat milk and 0.1% Triton X-100 for 1 h with orbital agitation. Hereupon, the membrane was incubated with the primary antibody overnight, at 4 °C, with agitation. The anti-BiP mouse monoclonal antibody was used at a working dilution of 1:1250, while the anti-GAPDH rabbit monoclonal antibody was used at a dilution of 1:10000.

A washing step was then performed with PBS 0.1% Triton X-100 for 30 min. Then, the secondary antibody was incubated for 2 h at room temperature. The anti-mouse secondary antibody was used at 1:2000, and the anti-rabbit secondary antibody at 1:3000. At this point, two more washing steps were performed with PBS 0.1% Triton X-100 for 15 min and then with PBS, again twice for 15 min. Finally, bands were detected by addition of a chemiluminescent substrate, WesternBright ECL HRP. The resulting bands were visualized on a ChemiDoc™ Imaging System (Bio-Rad, Hercules, CA, USA). Their relative optical density was calculated by densitometry resorting to the Fiji/ImageJ software version 1.51 (Fiji, Madison, WI, USA) and normalized against the loading control (GAPDH). Results are presented as mean ± SEM of, at least, three independent experiments.

### 2.9. Statistical analysis

The statistical analysis was performed using GraphPad Prism 8 software. We resorted to the *t*-test with a level of significance of *p <* 0.05.

## 3. Results and Discussion

### 3.1. Chemical space of the library

We compiled a library of 134 molecules of natural origin (**Table S2**). The average molecular weight was 358 g/mol, the inferior limit being 110.1 g/mol (catechol and *p*-hydroquinone) and the upper limit 1131.3 (β-escin) (**Fig. 1A**). The computed natural product-likeness score, a parameter that tries to quantify how much a given molecule is “natural product-like” [27] had an average of 1.29 (min: -0.4, max: 2.72, **Fig. 1A**). Over 50% of the molecules displayed no violations of Lipinski’s Rule of 5 with over 30% displaying 2 or more (**Fig. 1A**), a relevant trait for candidate drugs.

For chemical ontology, we have used the computed chemical classes generated by ClassyFire [28]. From a chemical point of view, the distribution of the molecules by chemical superclass can be found in **Fig. 1B**. Phenylpropanoids and polyketides corresponded to the majority of the entries (65.9%), followed by benzenoids (14.4%) and lipids or lipid-like molecules (6.1%). Within phenylpropanoids, flavonoids were the most prevalent molecules (**Fig. 1C**), followed by cinnamic acids and their derivatives. In the case of benzenoids, simple phenols were predominant, while in the case of lipids, prenol lipids were the most representative (**Fig. 1C**). The chemical structures of all of the molecules are represented in **Fig. S1**, separated according to the herein mentioned chemical superclasses.

### 3.2. Impact of molecules upon cell viability

Having performed the characterization of our in-house library of over 130 natural products, considering that new candidates for countering ER stress should be devoid of toxicity, we proceeded to the evaluation of the impact of all the molecules in the viability of fibroblasts, as assessed by the MTT reduction assay. Fibroblasts present a highly developed endoplasmic reticulum, since their function relates to the production of components of the extracellular matrix, and thus are particularly susceptible to ER stress, reason for which they were chosen as experimental model in this work [29].

Given the high number of molecules involved, all compounds were tested at a single concentration (50 µM). Statistically significant decreases in MTT reduction were interpreted as negative impact on cellular viability and the corresponding molecules excluded from subsequent experiments. As evidenced by **Fig. 2**, 37 molecules (depicted in **Table S2**) were dropped from the pipeline for their toxicity.

**Fig. 2.**
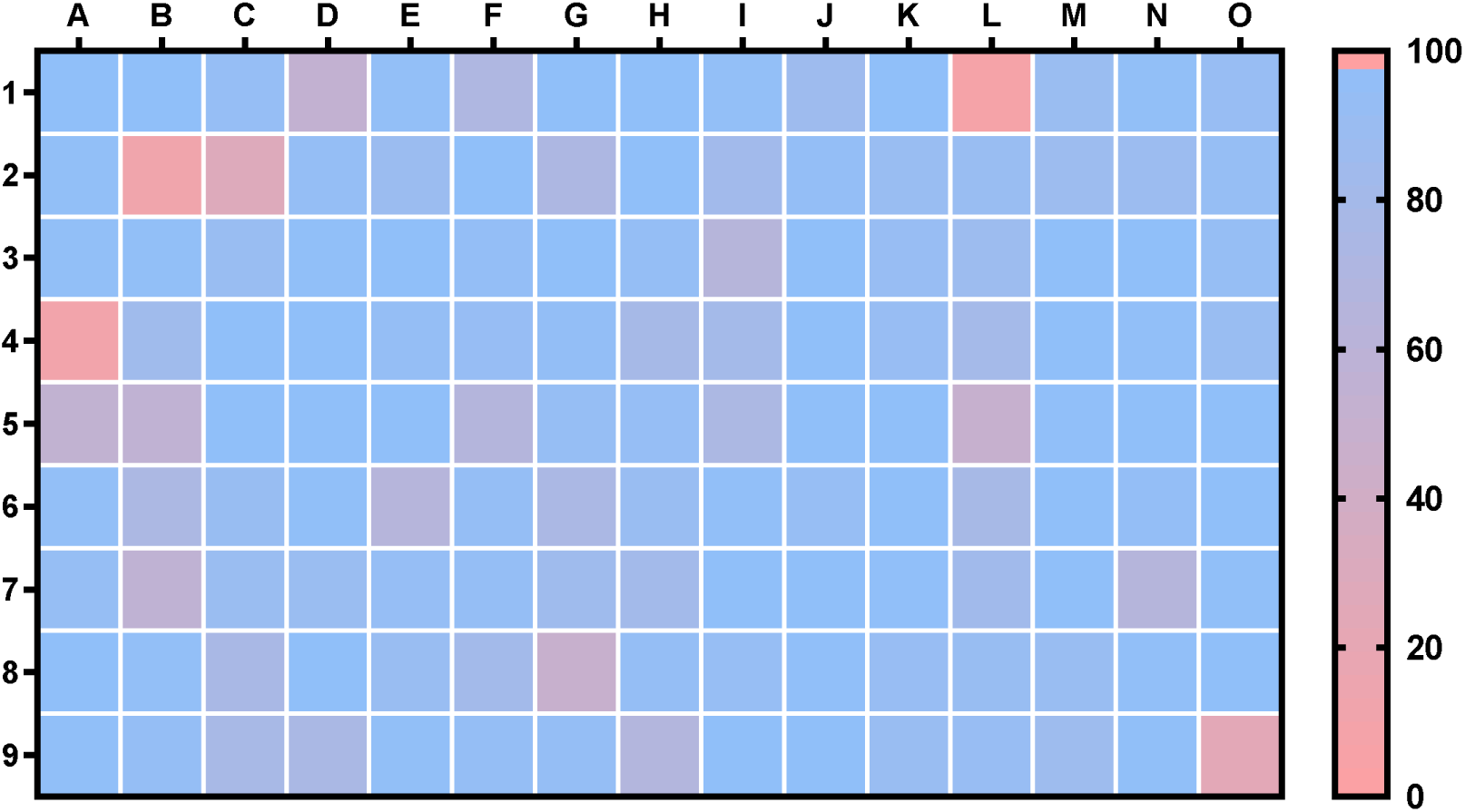
Impact of a library of natural products on MRC-5 fibroblasts viability, as determined by MTT reduction assays. Results represent the percentage of the control and correspond to, at least, three independent experiments, each performed in triplicate. [2-column]

The remaining 97 compounds were screened for potentially protective capacity against ER stress. The legend for the heatmap presented in **Fig. 2** can be consulted in **Table S2**, while **Fig. S2** contains the individual results for each molecule represented in the bar charts.

### 3.3. Effect of the library upon cytosolic calcium levels

The next step involved the screening of the nontoxic molecules for potential protection against the onset of ER stress. Towards this goal, and considering the high number of molecules still in the pipeline, we selected changes on calcium levels as an indicator of compromised ER homeostasis. The maintenance of the high concentration of calcium in the ER lumen creates an appropriate environment for protein folding, and thus changes on its levels heavily impact ER function [30, 31]. This organelle is able to maintain its calcium levels due to the abundance of resident calcium-buffering proteins, like calnexin and calreticulin [32]. Under stress conditions, the ER may release calcium through channels, such as IP_3_R and RyR. Eventually, this may lead to increased calcium in mitochondria, indirectly inducing the generation of ROS and triggering the dissipation of the mitochondrial membrane potential, ultimately leading to the release of resident proapoptotic factors [33].

To evaluate the potential protective effect of the molecules under study, we conducted experiments with the fluorescent probe Fura-2/AM. Calcium ions were detected employing the divalent cation ionophore A23187 (5 µM) as a positive control for calcium leakage into the cytosol and molecules that could inhibit or counter its effect were searched. Calcium ionophore A23187, or calcimycin, is a calcium-binding molecule that increases the cellular permeability of ionic calcium [34]. There are multiple examples in the literature of the use of this molecule to increase cytosolic calcium levels, inducing ER stress, UPR activation and eventual apoptosis or autophagy [35–37].

Under these experimental conditions, we were able to identify several molecules that significantly prevented or ameliorated the calcium overload caused by A23187, as displayed in **Fig. 3**. Such molecules included six polyphenols (5-deoxykaempferol, delphinidin, fisetin, naringenin, naringin, quercetin-3-*O*-β-D-glucoside), one anthraquinone (sennoside B), one fatty acid (myristic acid) and one biogenic polyamine (spermine). The full list of molecules tested in this assay can be consulted in **Table S3** while **Fig. S3** represents these data in the bar charts.

**Fig. 3.**
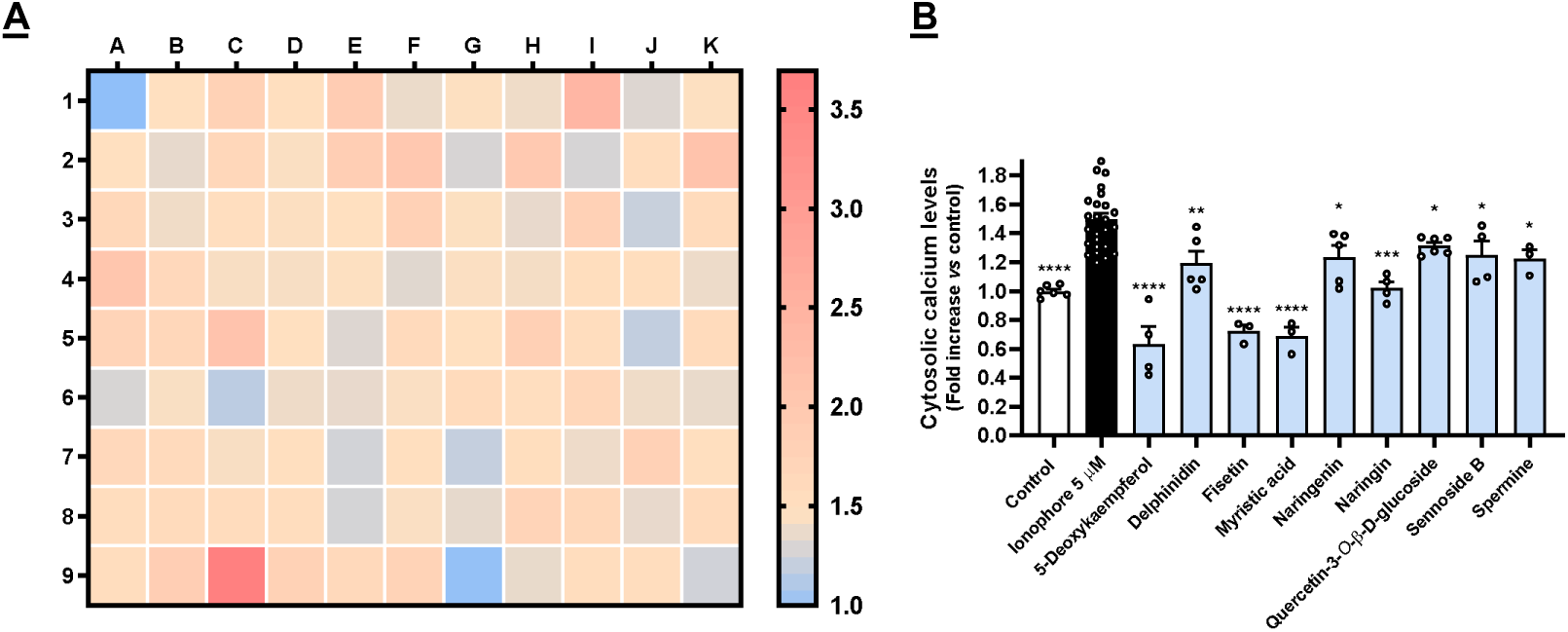
Cytosolic calcium levels in MRC-5 fibroblasts in response to co-incubation of calcium ionophore A23187 at 5 µM and nontoxic molecules (50 µM), as determined by the fluorescence of the probe Fura-2/AM. **A**) Heatmap representing all molecules tested; **B**) Molecules that caused a statistically significant decrease in calcium levels when compared to the positive control. Results correspond to the mean of, at least, three independent experiments, each performed in triplicate. [2-column]

### 3.4. ER stress counter candidates inhibit thapsigargin-induced UPR signaling in MRC-5 fibroblasts

The non-toxic molecules that successfully lowered Ca^2+^ levels were considered potential candidates for countering ER stress. As so, they were evaluated for their effect upon UPR signaling pathways.

Thapsigargin, a natural sesquiterpene lactone that is widely used as UPR inducer, was the positive control. This molecule acts as an irreversible SERCA pump inhibitor, leading to an impaired calcium homeostasis and subsequent onset of organized cell death mechanisms [2]. Accordingly, as evidenced by **Fig. 4**, we can see that thapsigargin (3 µM) significantly increased the expression of all the genes of interest, namely the transcription factors *DDIT3* and *ATF4*, the chaperones *HSPA5* and *HSP90B1* (BiP and GRP94 proteins, respectively), and *EDEM1*, a gene coding for the ER degradation-enhancing alpha-mannosidase-like protein 1, an enzyme involved in ER-associated degradation.

**Fig. 4.**
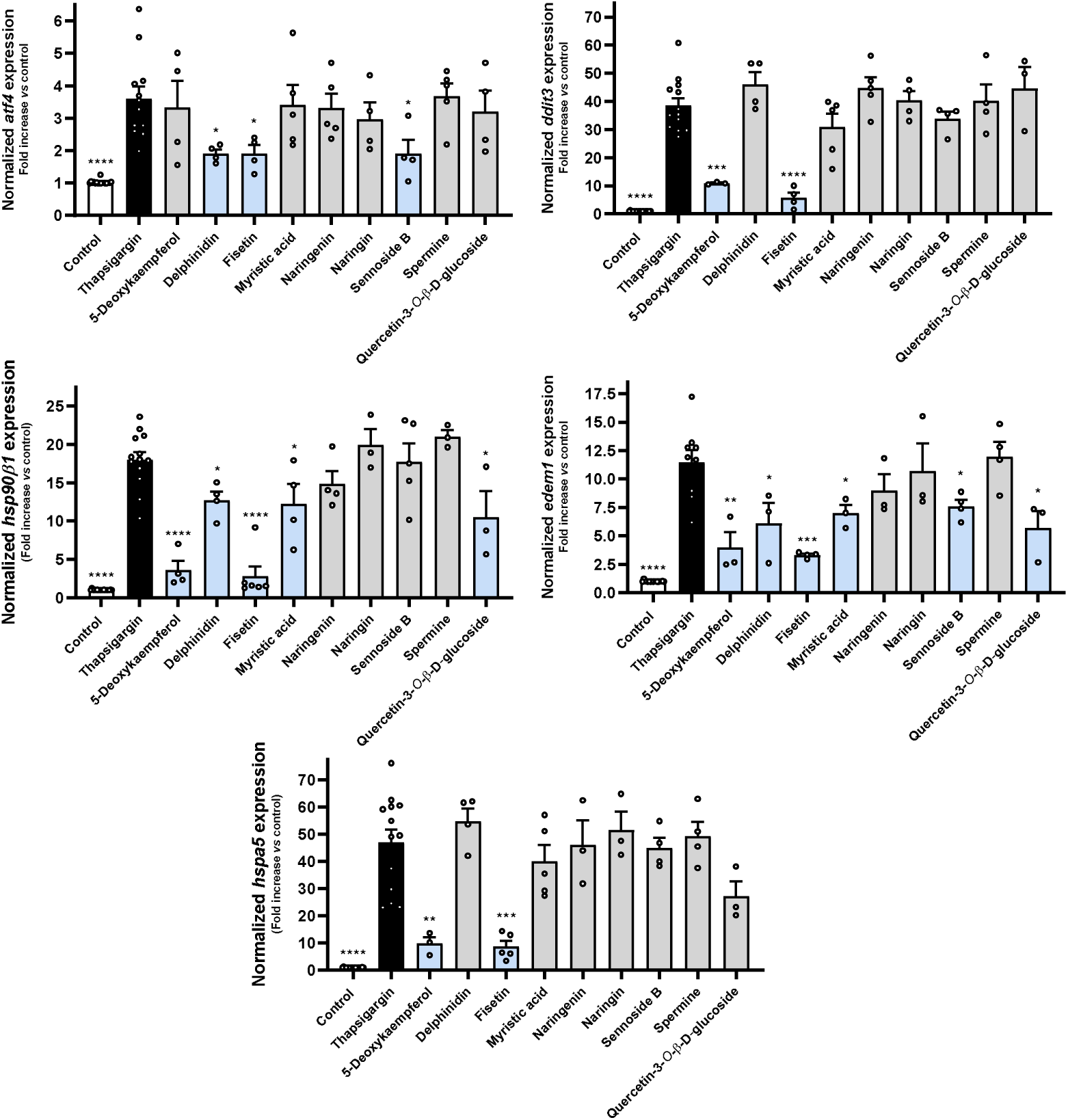
Effect of 5-deoxykaempferol, delphinidin, fisetin, myristic acid, naringenin, naringin, sennoside B, spermine and quercetin-3-*O*-β-D-glucoside on Tg-induced UPR signaling, as determined by RT-qPCR. Results correspond to the mean ± standard error of the mean of, at least, four independent experiments, each performed in duplicate. *GAPDH* was used as a reference gene for normalization of expression. [2-column]

5-Deoxykaempferol at 50 µM inhibited the expression of *DDIT3*, *HSP90B1*, *EDEM1* and *HSPA5*, even though it did not significantly decrease *ATF4* expression. Fisetin was the only molecule to prevent the increase of all target genes in a significant manner. Delphinidin negatively impacted the expression of *ATF4*, *HSP90B1* and *EDEM1*. Myristic acid and quercetin-3-*O*-β-D-glucoside displayed a similar pattern in the protection against UPR activation, by modulating *HSP90B1* and *EDEM1* expressions. Sennoside B acted upon *ATF4* and *EDEM1*, while naringenin, naringin and spermine failed to inhibit any of the selected genes.

In light of these results, we show that 6 out of the 9 molecules that effectively countered the calcium overload were, simultaneously, capable of downregulating UPR-associated genes, which suggests that the assessment of cytosolic calcium levels is a reliable method to screen of the capacity to counter ER stress in this experimental model. Considering that 5-deoxykaempferol, delphinidin, fisetin, myristic acid, sennoside B and quercetin-3-*O*-β-D-glucoside were capable of significantly preventing the Tg-induced increased expression of, at least, two UPR-related genes, they were selected for subsequent studies. In contrast, naringin, naringenin and spermine failed to inhibit the expression of any of the analyzed genes, thus they were dropped from the pipeline.

### 3.5. Molecules that counter ER stress ameliorate protein aggregation in MRC-5 fibroblasts and attenuate UPR activation at the protein level

As discussed earlier, activation of the UPR affects the capacity of cells to properly fold proteins, which frequently results in the formation of protein aggregates and may lead to proteinopathies [38, 39]. Conversely, several diseases, notably those of neurodegenerative nature, are associated with these aggregates [40]. As so, new molecules capable to countering ER stress are expected to be able to prevent or lower the amounts of protein aggregates in a biological context. To confirm this, we proceeded to an experimental model of ER stress-triggered protein aggregation using Aβ_25-35_ as a protein aggregation inducer. Aβ_25-35_ is an active fragment of Aβ_42_ [41], a peptide that is associated to the pathogenesis of Alzheimer’s disease (AD) and has been characterized in experimental models, where it can lead to the formation of protein aggregates *in vitro* and AD *in vivo* [42].

As shown in **Fig. 5A**, incubation of MRC-5 cells with Aβ_25-35_ elicits protein aggregation, as assessed with thioflavin T, a benzothiazole that presents high fluorescence when bound to β-sheet-rich structures common in protein aggregates, both in a microplate-based (**Fig. 5A**) and an immunocytochemistry method (**Fig. 5B**). In the same conditions, delphinidin, fisetin, myristic acid, sennoside B and quercetin-3-*O*-β-D-glucoside significantly inhibited this effect. These results show that the molecules that had been selected on the basis of their ability to attenuate UPR pathways were able to act upon the endpoint biological effect of this cascade, namely protein aggregation.

**Fig. 5.**
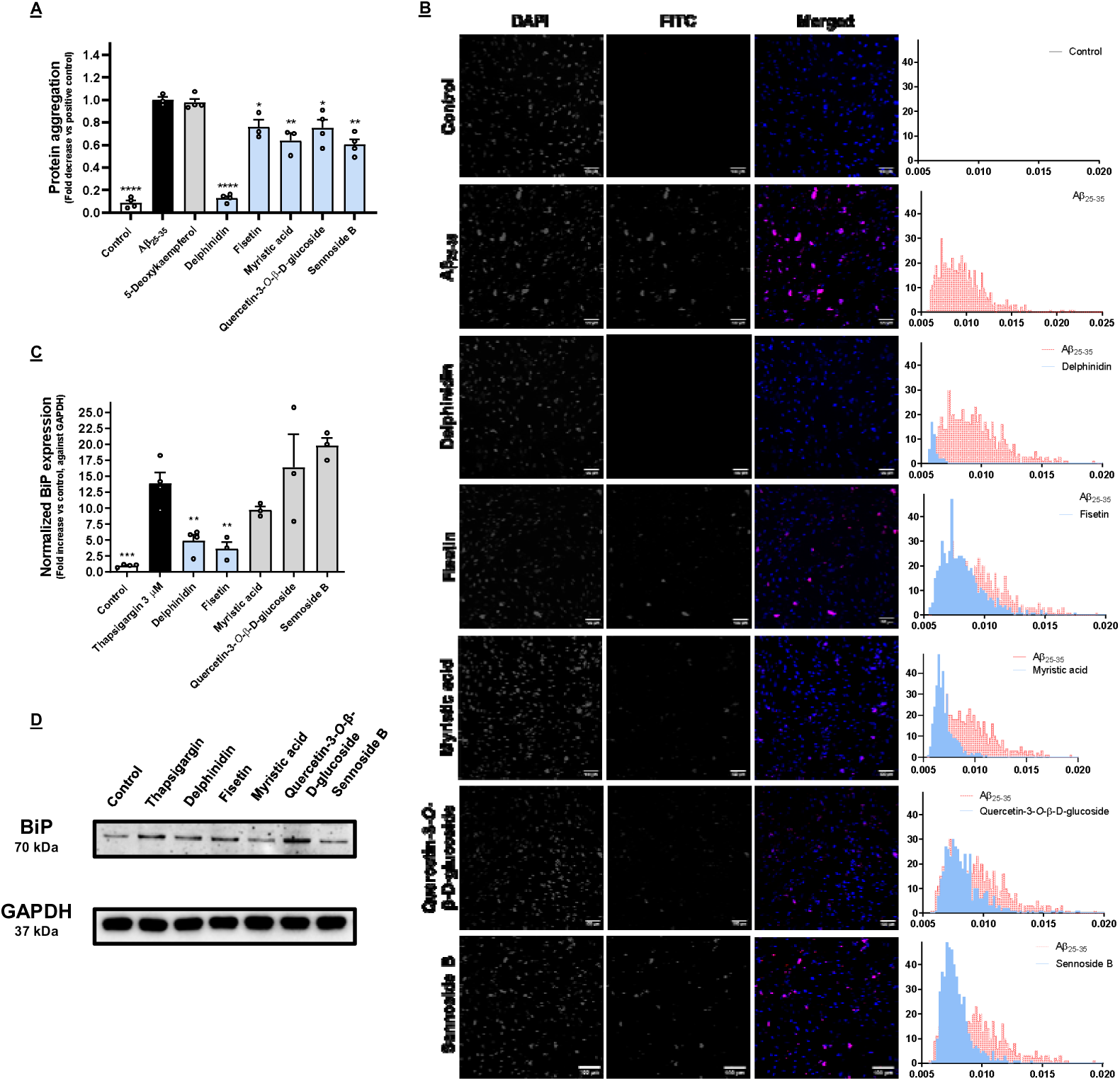
**A:** Impact of delphinidin, fisetin, myristic acid, sennoside B and quercetin-3-*O*-β-D-glucoside on Aβ_25-35_-induced protein aggregation in MRC-5 cells, determined by the fluorescence of thioflavin T. Results correspond to the mean ± standard error of the mean of, at least, three independent experiments, individually performed in triplicate. **B:** The protective effect of bioactive molecules was captured under a fluorescence microscope. The fluorescent signal of thioflavin T is represented in the histograms. **C:** Effect of delphinidin on Tg-induced BiP expression in MRC-5 cells, as evaluated by Western blotting. Results correspond to the mean ± standard error of the mean of, at least, three independent experiments. **D:** Images obtained through Western blotting analysis. [2-column]

As mentioned before, the UPR relies on three major signaling branches. They are all governed by the major chaperone BiP, coded by the gene *HSPA5*. Under homeostatic conditions, BiP remains bound to the three major UPR players (PERK, IRE1 and ATF6), releasing them whenever it recognizes the presence of misfolded proteins. This extends the permanence of these proteins in the ER lumen, in order to promote their correct processing. For these reasons, BiP is a key regulator of UPR signaling and its activation prevents protein aggregation [43]. Furthermore, it is described that its expression is increased throughout the course of neurodegenerative disease [44]. Our Western blotting results show that the exposure of Tg-insulted MRC-5 cells to delphinidin and fisetin clearly reduced BiP expression, and, consequently, we presume that the UPR activation in the presence of these molecules is mitigated.

### 3.6. Selected molecules counter protein aggregation and UPR activation at the translational level in neuronal cells

Given the promising results on MRC-5 fibroblasts, we repeated the experiment, this time with SH-SY5Y cells, in order to analyze whether these compounds could also be protective on this neuronal cell line. The SH-SY5Y cell line has been widely used and described as a good *in vitro* model for the study of protein aggregation [45–48], a process that has been shown to be both consequence and cause of misfolded proteins in the ER and subsequent activation of the UPR [49].

Relevantly, in the case of AD it has been shown that increased levels of phosphorylated PERK and IRE1α are found in the hippocampus of the patients and that they colocalize with phosphorylated tau. Furthermore, treatment of cells with Aβ_42_, the major component of amyloid plaques, induces CHOP expression, which is a product of the PERK and ATF6 branches of the UPR [44, 50, 51]. For this assay, the concentrations of the molecules under study had to be adjusted to the highest nontoxic concentration towards this cell line. Fisetin was tested at 25 µM, while the remaining molecules were tested at the concentrations used in MRC-5 cells (50 µM). The effect of this compound was not replicated on SH-SY5Y cells. We hypothesize that this may be due to the reduction of the concentration that was necessary in order not to impact cell viability in this model. On the other hand, the effect of delphinidin translates to these cells, virtually nullifying the advent of protein aggregates, as evidenced by **Fig. 6A** and **B**.

**Fig. 6.**
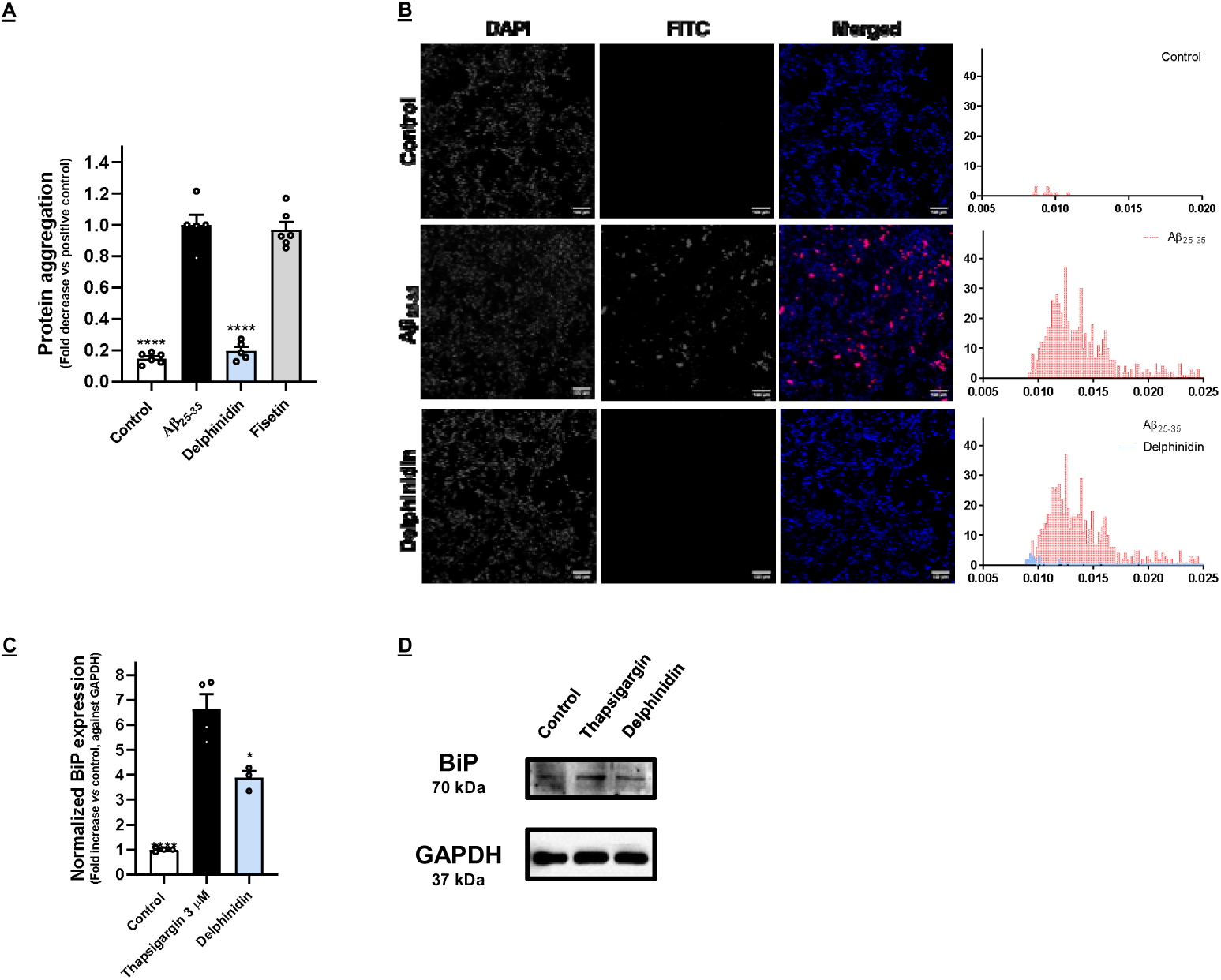
**A:** Effect of delphinidin and fisetin on Aβ_25-35_-induced protein aggregation in SH-SY5Y cells, as evaluated by determination of the fluorescence of thioflavin T. Results correspond to the mean ± standard error of the mean of, at least, three independent experiments, each performed in triplicate. **B:** The effect of bioactive molecules was observed under a fluorescence microscope. The fluorescent signal of thioflavin T is represented in the histograms. **C:** Effect of delphinidin on Tg-induced BiP expression in SH-SY5Y cells, as evaluated by Western blotting. Results correspond to the mean ± standard error of the mean of, at least, three independent experiments. **D:** Example of the obtained Western blotting results. [2-column]

### 3.7. Defining a chemical space for future drug development

Considering all the information generated during this work, together with the chemical, structural and topographical information available for the molecules under study, we were interested in understanding if the physico-chemical properties of the library could, *per se*, be a good indicator for its activity. To this end, we took all the molecules used and computed multiple pairs of their properties, sorting them as active or inactive according to the results of the aggregation assays depicted in **Fig. 5**. While there seemed to be a tendency of active molecule to be apart from inactive molecules in some cases (**Fig. S4**), it soon became apparent that a pairwise comparison could leave behind important features of the molecules, as it reduces high dimensional data to just two dimensions. As so, we have decided to use other methodologies that could take into account all the features of the library, namely dimensionality reduction algorithms, which allows the combination of different features into principal components that can be more easily graphed and understood. We first conducted a correlation study in order to identify molecular features that were highly correlated, which could negatively impact the performance of the statistical analysis given the redundancy it would add. **Fig. 7** shows the original features, where it is clear that many are highly correlated, for example, the features “total atom number” and “number of carbons” (0.98), as one directly affects the other. Such features were dropped, and the remaining ones were used. We also decided to assess the robustness of the methods in identifying the chemical space occupied by the molecule that was shown to be active in both cell lines, namely delphinidin.

**Fig. 7.**
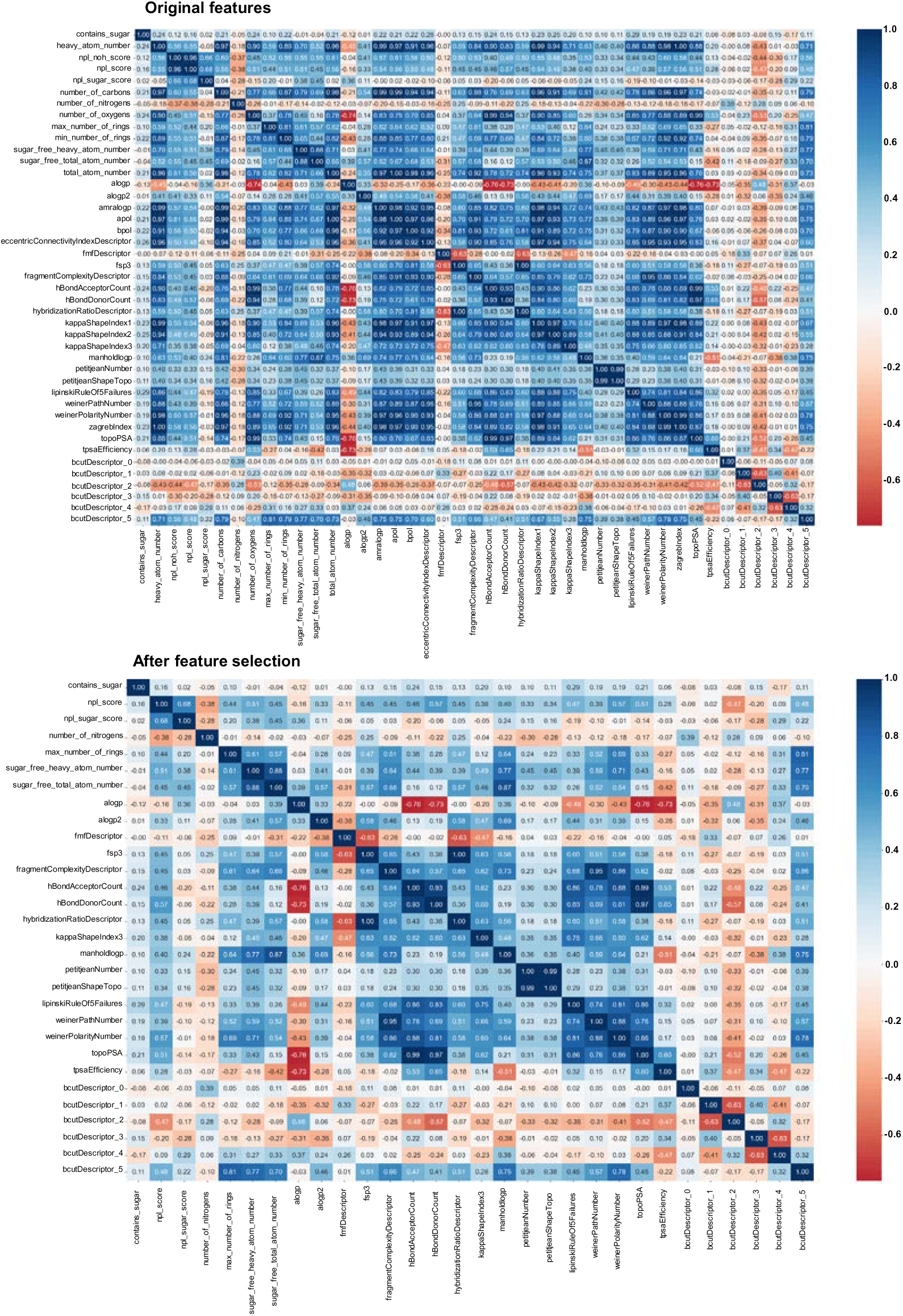
Correlation matrix for the features used for the chemical library used in this work. [2-column]

We initially used the classical algorithm for principal component analysis (PCA), which is widely used both for chemometric and biological data [52, 53]. As shown, PCA was unable to identify a chemical space occupied solely by the molecule that was shown to be active in all the assays used (**Fig. 8A**), the same being true for multidimensional scaling (MDS, **Fig. 8B**). In light of this, we applied a suite of other algorithms, both individually and in tandem with PCA, namely uniform manifold approximation and projection (UMAP) and t-distributed stochastic neighbor embedding (t-SNE).

**Fig. 8.**
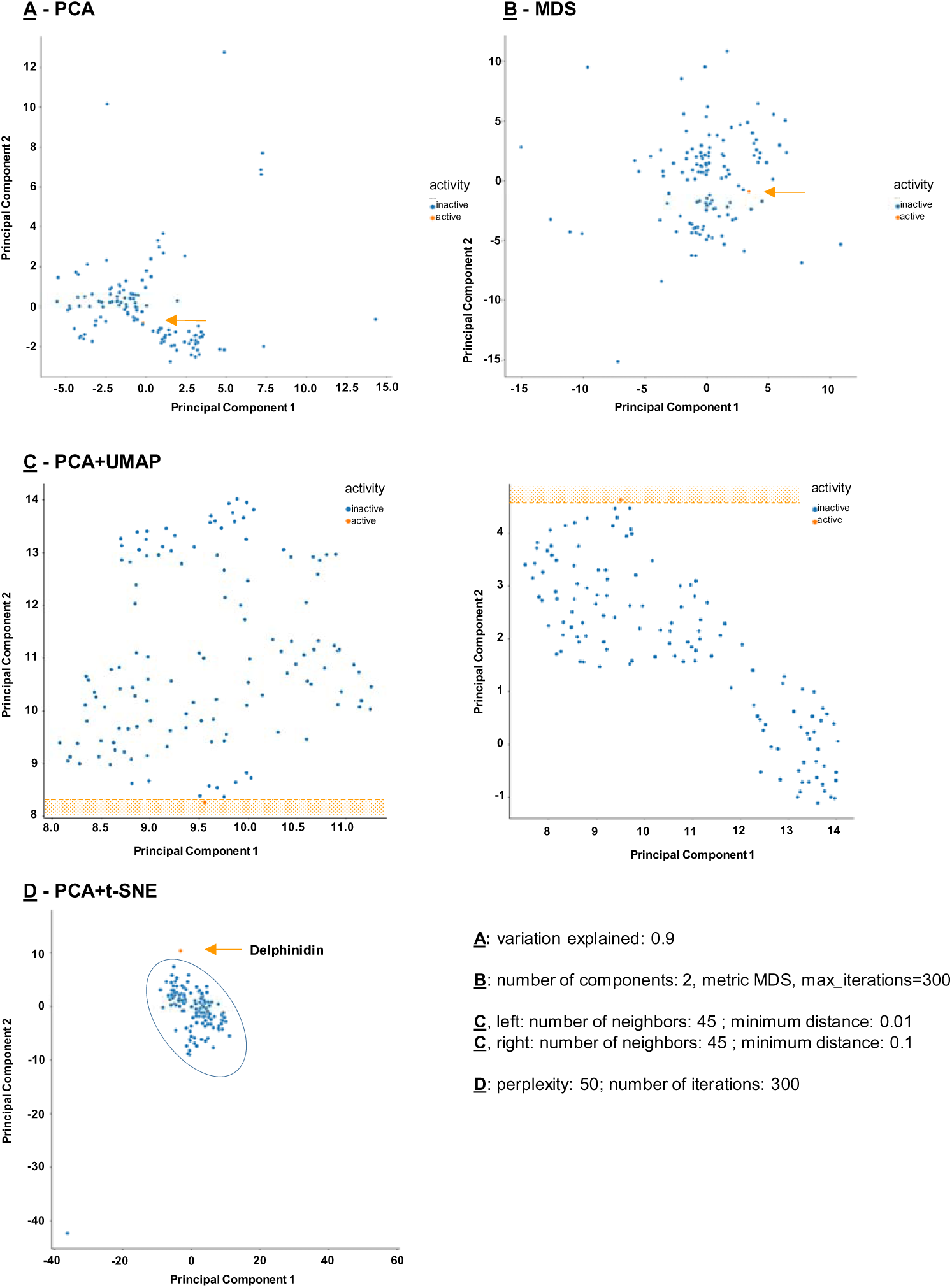
Different dimensionality reduction methods applied to chemical library under study. The active molecule delphinidin is presented in orange and inactive molecules in blue. [2-column]

Differently from PCA and MSD, the combined used of PCA+UMAP led to a 2-principal component-based space, in which delphinidin clearly occupies an outlier space. We have tried distinct combinations of parameters, as stated in the Materials section, the number of neighbors of 45 giving the best performance, both for minimum distance of 0.01 and 0.1. In order to confirm this result, we have employed an additional algorithm, namely PCA+t-SNE. Again, it was shown that delphinidin, together with an outlier, occupies a space that is distinct from over 98% of the library.

Taken together, these results show that chemometric tools, in tandem with biological data generated from significant number of samples, can be used for future selection of drug candidates relying on data science-based approaches. As so, the composition of the principal components identified can be used for the subsequent selection of novel molecules capable of countering ER stress by screening large databases of molecules for ideal chemical descriptors.

## 4. Conclusions

Our framework blending data science-based methods to biological data resulted in the identification of several potentially active molecules against the onset of ER stress. Delphinidin stood out as the most promising candidate against UPR activation and protein aggregation in a cellular model of neurodegenerative disease, being capable to simultaneously modulate calcium levels and downregulate UPR pathways with meaningful biological endpoints.

In light of our results, it is safe to assume that the natural product chemical space may provide molecules that ultimately lead to the development of novel pharmacological strategies to counter ER stress-related pathologies, as is the case of neurodegenerative diseases.

## 5. Acknowledgements

The work was supported by UIDB/50006/2020 with funding from FCT/MCTES through national funds. Daniela Correia da Silva extends special thanks to the Fundação para Ciência e Tecnologia (FCT) for the grant (SFRH/BD/130998/2017). Authors would also like to acknowledge BioRender.com as a platform for image creation.

