## Supplementary tables and Figures for "A pipeline for natural small molecule inhibitors of endoplasmic reticulum stress": Supplementary Material_UPR_Pharmacological Research.docx

**Table S1.** Variables used in the chemometric analysis.

| **Variable** | **Description** |
| --- | --- |
| coconut_id | Identifier used in the COCONUT database |
| contains_sugar | Bolean for the presence of sugars |
| heavy_atom_number | Number of heavy atoms |
| name | Name of the molecule |
| contains_ring_sugars | Bolean for the presence of sugars |
| molecular_formula | Molecular formula |
| molecular_weight | Molecular weight |
| npl_noh_score | Natural product-likeness score without taking into account any hydrogen in the molecular structure |
| npl_score | Natural product-likeness score |
| npl_sugar_score | NP-likeness score computed on the glycosylated molecule |
| number_of_carbons | Number of carbons |
| number_of_nitrogens | Number of nitrogens |
| number_of_oxygens | Number of oxygens |
| max_number_of_rings | Maximal number of rings |
| min_number_of_rings | Minimal number of rings |
| sugar_free_heavy_atom_number | Heavy atoms number, excluding sugars |
| sugar_free_total_atom_number | Total atom number, excluding sugars |
| total_atom_number | Total atom number |
| alogp | AlogP |
| alogp2 | AlogP squared |
| amralogp | AMR - molar refractivity |
| apol | Sum of the atomic polarizabilities |
| bpol | BPol descriptor |
| eccentricConnectivityIndexDescriptor | Eccentric Connectivity Index Descriptor |
| fmfDescriptor | fmf Descriptor |
| fsp3 | fsp3 |
| fragmentComplexityDescriptor | Fragment Complexity Descriptor |
| hBondAcceptorCount | Hydrogen bond acceptor count |
| hBondDonorCount | Hydrogen bond donor count |
| hybridizationRatioDescriptor | Hybridization Ratio Descriptor (fraction of sp3 carbons to sp2 carbons) |
| kappaShapeIndex1 | First kappa shape index |
| kappaShapeIndex2 | Second kappa shape index |
| kappaShapeIndex3 | Third kappa shape index |
| manholdlogp | LogP descriptor (Mannhold version) |
| petitjeanNumber | Petit Jean Number |
| petitjeanShapeTopo | Petitjean geometrical shape index |
| lipinskiRuleOf5Failures | Number of Lipinski Rule of 5 violations |
| vertexAdjMagnitude | Vertex adjacency information |
| weinerPathNumber | Wiener Path Number |
| weinerPolarityNumber | Wiener Polarity Number |
| zagrebIndex | Zagreg Index |
| topoPSA | Topological polar surface area descriptor |
| tpsaEfficiency | Fractional polar surface area descriptor |
| iupac_name | IUPAC name |
| chemicalClass | Chemical Class |
| chemicalSubClass | Chemical SubClass |
| chemicalSuperClass | Chemical SuperClass |
| directParentClassification | Classification of parent molecule |

**Table S2.** Molecules tested in the cell viability assays. Molecules in red were excluded given their toxicity.

|  | **A** | **B** | **C** | **D** | **E** |
| --- | --- | --- | --- | --- | --- |
| **1** | Control | 5,7,8-Trihydroxyflavone | Betanin | Chrysophanol | Ellagic acid |
| **2** | (-)-Norepinephrine | 5,8-Dihydroxy-1,4-naphtoquinone | Betulin | Cinnamic acid | Emodin |
| **3** | (+/-)-Dihydrokaempferol | 5-Deoxykaempferol | Boldine | Coumarin | Eriocitrin |
| **4** | 1,4-Naphtoquinone | 5-Methoxypsoralen | Caffeine | Cyanidin | Eriodictyol |
| **5** | 18α-Glycyrrhetinic acid | Acacetin | Catechol | Cynarin | Eriodictyol-7-*O*-glucoside |
| **6** | 3,4-Dihydrobenzoic acid | Ajmalicine | Chlorogenic acid | Daidzein | Eupatorin |
| **7** | 3,4-Dimethoxycinnamic acid | Apigenin | Cholesta-3,5-diene | Delphinidin | Ferulic acid |
| **8** | 3-Hydroxybenzoic acid | Apigetrin | Chrysin | Diosmetin | Fisetin |
| **9** | 4-Hydroxybenzoic acid | Berberine | Chrysoeriol | Diosmin | Flavanone |
|  | **F** | **G** | **H** | **I** | **J** |
| **1** | Galangin | Hesperetin | Isorhamnetin-3-*O*-rutinoside | Liquiritigenin | Mangiferin |
| **2** | Galanthamine | Hesperidin | Isorhoifolin | Lupeol | Maritimein |
| **3** | Gallic acid | Homoeriodictyol | Juglone | Lutein | Myricetin |
| **4** | Genistein | Homoorientin | Kaempferide | Luteolin | Myricitrin |
| **5** | Genkwanin | Homovannilic acid | Kaempferol | Luteolin tetramethylether | Myristic acid |
| **6** | Gentisic acid | Hydroquinone | Kaempferol-3-*O*-rutinoside | Luteolin-3’,7-di*-O*-glucoside | Myrtillin |
| **7** | Guaiaverin | Hyoscyamine | Kaempferol-7-*O*-glucoside | Luteolin-4'-*O*-glucoside | Naringenin |
| **8** | Guaiazulene | Isorhamnetin | Kaempferol-7-*O*-neohesperidoside | Luteolin-7-*O*-glucoside | Naringenin-7-glucoside |
| **9** | Herniarin | Isorhamnetin-3-*O*-glucoside | Linarin | Malvidin | Naringin |
|  | **K** | **L** | **M** | **N** | **O** |
| **1** | Narirutin | Plumbagin | Quercetin-3-*O*-*ß*-*D*-glucoside | Sennoside B | Tiliroside |
| **2** | Oleuropein | Pyrogallol | Quercitrin | Silibinin | Trigonelline |
| **3** | Orientin | Quercetagetin | Rhoifolin | Spermine | Vanillin |
| **4** | *p*-Coumaric acid | Quercetin | Robinin | Sulfuretin | Verbascoside |
| **5** | Pelargonidin | Quercetin-3,3’,4',7-tetramethylether | Rosmarinic acid | Swertiamarin | Vicenin-2 |
| **6** | Pelargonin | Quercetin-3,4'-dimethylether | Rutin | Taxifolin | Vitexin |
| **7** | Phloroglucinol | Quercetin-3-methylether | Saponarin | Tectochrysin | Vitexin-2-*O*-rhamnoside |
| **8** | Pinocembrin | Quercetin-3-*O*-(-6-acetylglucoside) | Scopolamine | Theobromine | Xanthone |
| **9** | Pyrogallol | Quercetin-3-*O*-glucuronide | Sennoside A | Theophylline | ß-escin |

**Table S3.** Molecules screened for their ability to preserve cytosolic calcium levels. Molecules in blue were selected for subsequent studies.

|  | **A** | **B** | **C** | **D** | **E** |
| --- | --- | --- | --- | --- | --- |
| **1** | Control | 5-Deoxykaempferol | Cinnamic acid | Eriocitrin | Gentisic acid |
| **2** | Ionophore | Apigetrin | Coumarin | Eriodictyol | Guaiaverin |
| **3** | (-)-Norepinephrine | Berberine | Cyanidin | Eriodictyol-7-*O*-glucoside | Herniarin |
| **4** | (+/-)-Dihydrokaempferol | Betanin | Cynarin | Ferulic acid | Hesperetin |
| **5** | 3,4-Dihydrobenzoic acid | Boldine | Daidzein | Fisetin | Homoeriodictyol |
| **6** | 3,4-Dimethoxycinnamic acid | Caffeine | Delphinidin | Flavanone | Homoorientin |
| **7** | 3-Hydroxybenzoic acid | Catechol | Diosmetin | Galanthamine | Homovannilic acid |
| **8** | 4-Hydroxybenzoic acid | Chlorogenic acid | Ellagic acid | Gallic acid | Isorhamnetin-3-*O*-glucoside |
| **9** | 5,7,8-Trihydroxyflavone | Cholesta-3,5-diene | Emodin | Genistein | Isorhamnetin-3-*O*-rutinoside |
|  | **F** | **G** | **H** | **I** | **J** |
| **1** | Isorhoifolin | Malvidin | Narirutin | Pyrogallol | Saponarin |
| **2** | Juglone | Maritimein | Oleuropein | Quercetin-3-*O*-ß-D-glucoside | Scopolamine |
| **3** | Kaempferol | Myricetin | Orientin | Quercetin-3-*O*-(-6-acetylglucoside) | Sennoside B |
| **4** | Kaempferol-3-*O*-rutinoside | Myricitrin | *p*-Coumaric acid | Quercetin-3-*O*-glucuronide | Silibinin |
| **5** | Kaempferol-7-*O*-neohesperidoside | Myristic acid | Pelargonidin | Quercitrin | Spermine |
| **6** | Liquiritigenin | Myrtillin | Pelargonin | Rhoifolin | Sulfuretin |
| **7** | Luteolin-3’,7-di-*O*-glucoside | Naringenin | Phloridzin | Robinin | Swertiamarin |
| **8** | Luteolin-4'-*O*-glucoside | Naringenin-7-glucoside | Phloroglucinol | Rosmarinic acid | Taxifolin |
| **9** | Luteolin-7-*O*-glucoside | Naringin | Pinocembrin | Rutin | Theobromine |
|  | **K** |  |  |  |  |
| **1** | Theophylline |  |  |  |  |
| **2** | Tiliroside |  |  |  |  |
| **3** | Trigonelline |  |  |  |  |
| **4** | Vanillin |  |  |  |  |
| **5** | Verbascoside |  |  |  |  |
| **6** | Vicenin-2 |  |  |  |  |
| **7** | Vitexin |  |  |  |  |
| **8** | Vitexin-2-*O*-rhamnoside |  |  |  |  |
| **9** | Xanthone |  |  |  |  |

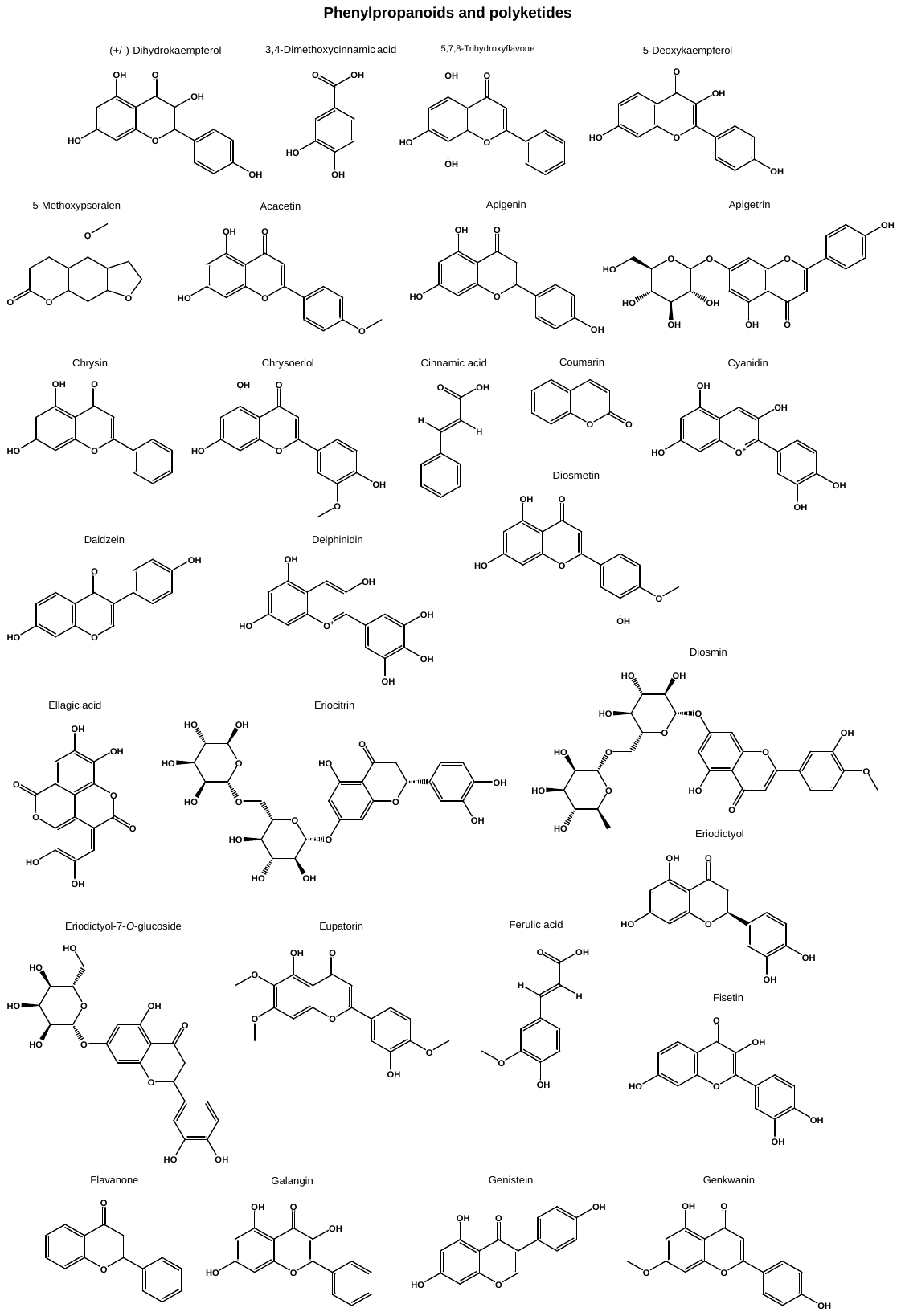

**Fig. S1**: Chemical structures of all tested molecules, grouped according to the respective chemical superclass according to Classyfire.

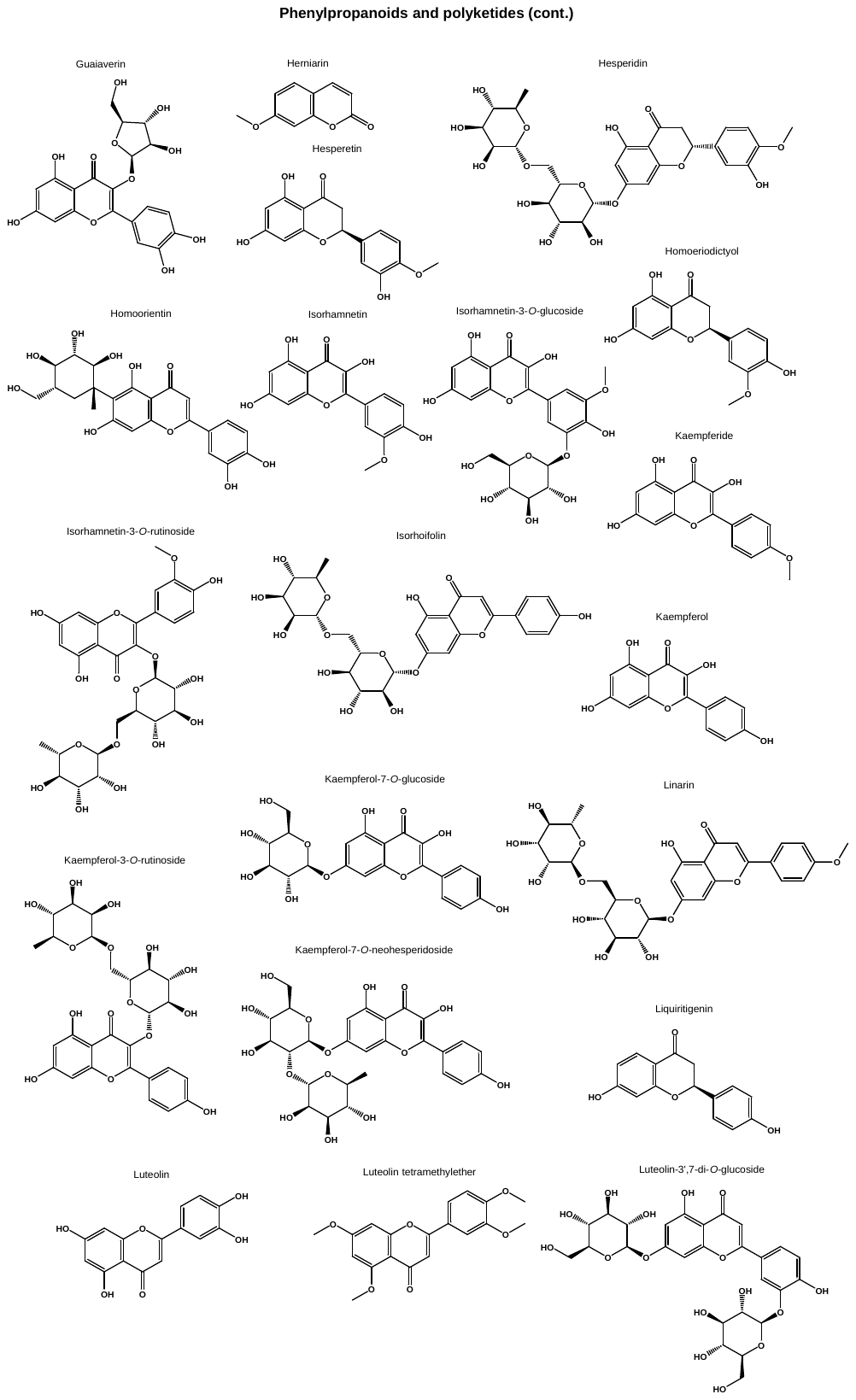

**Fig. S1 (cont.)**: Chemical structures of all tested molecules, grouped according to the respective chemical superclass according to Classyfire.

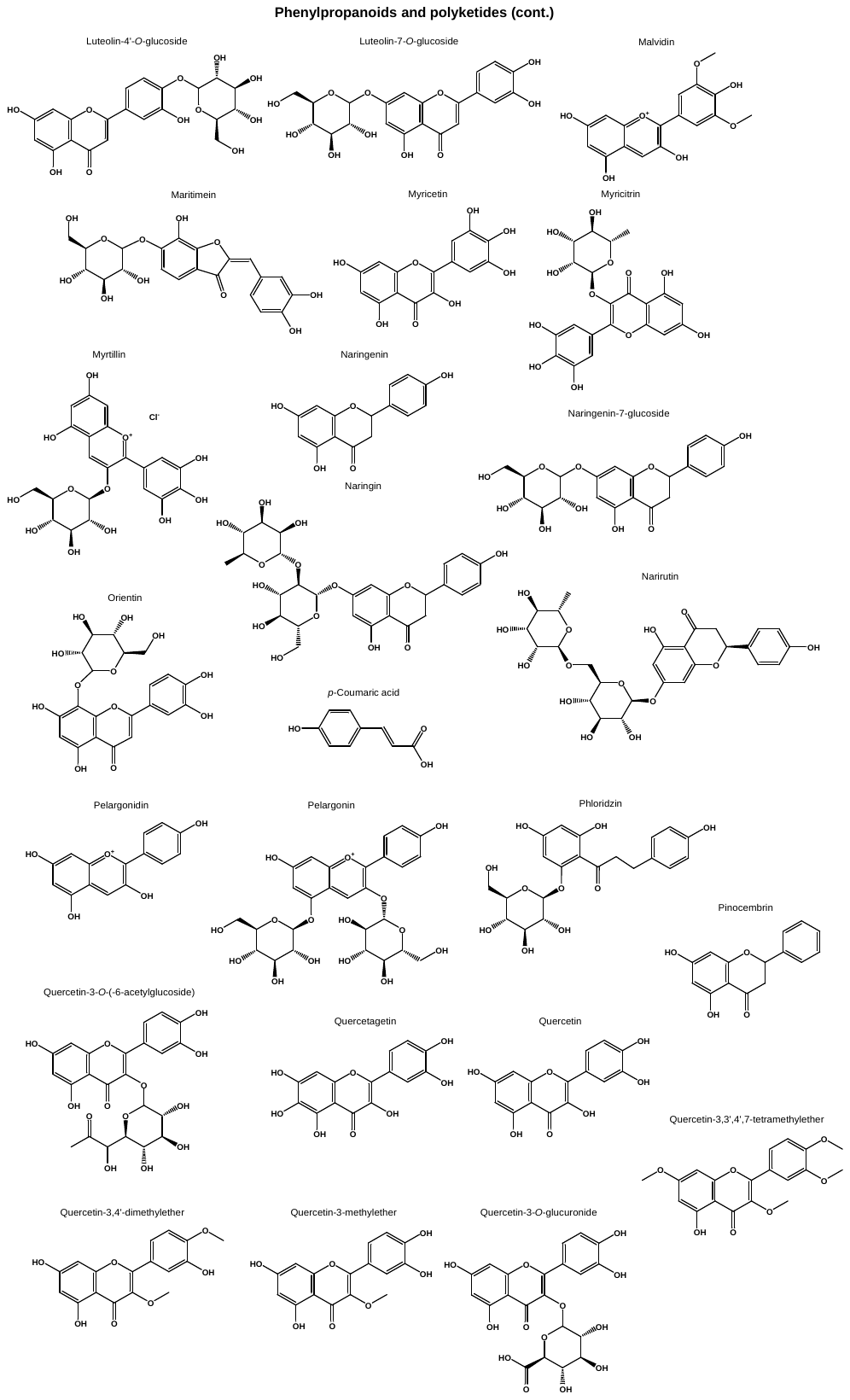

**Fig. S1 (cont.)**: Chemical structures of all tested molecules, grouped according to the respective chemical superclass according to Classyfire.

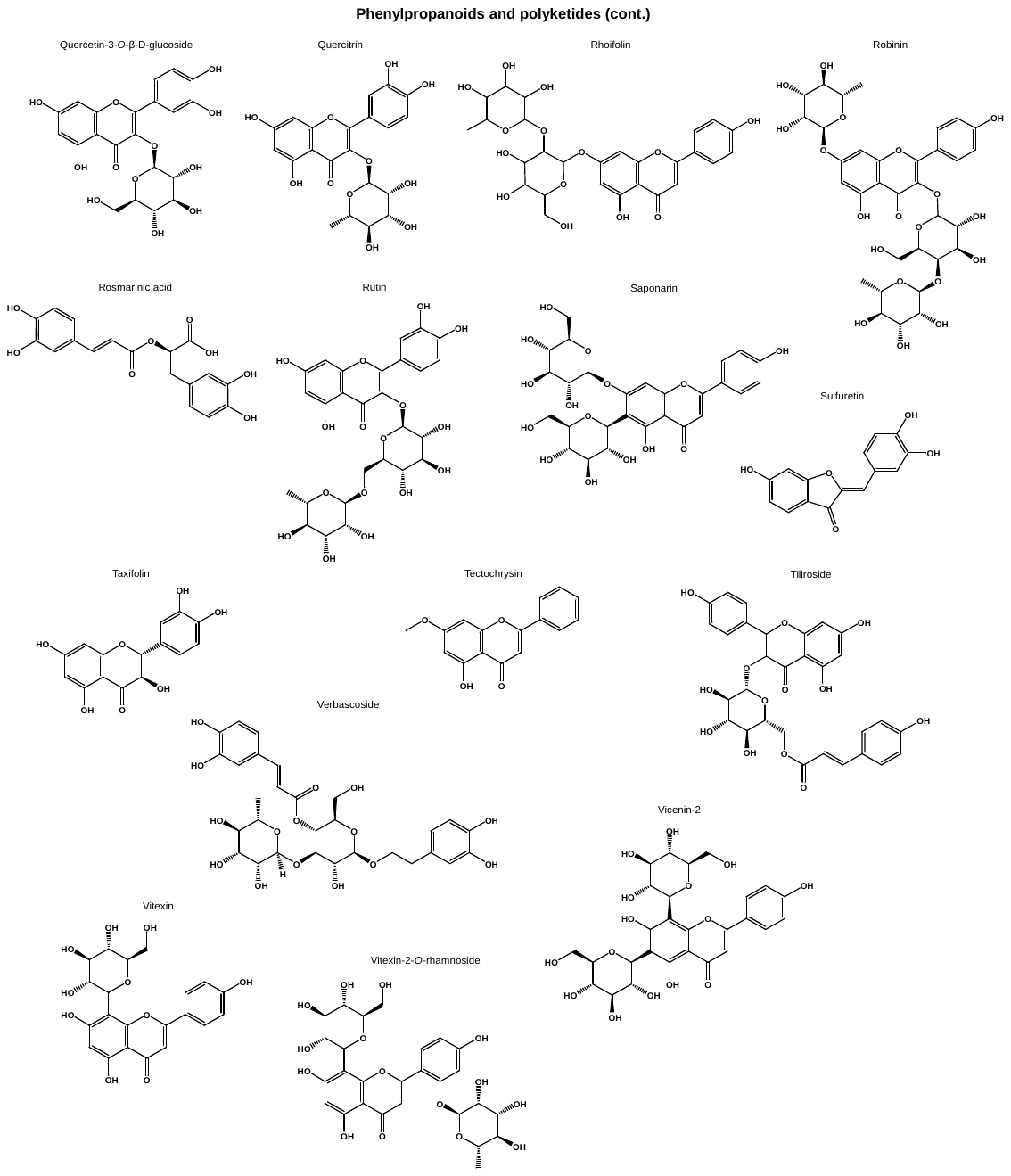

**Fig. S1 (cont.)**: Chemical structures of all tested molecules, grouped according to the respective chemical superclass according to Classyfire.

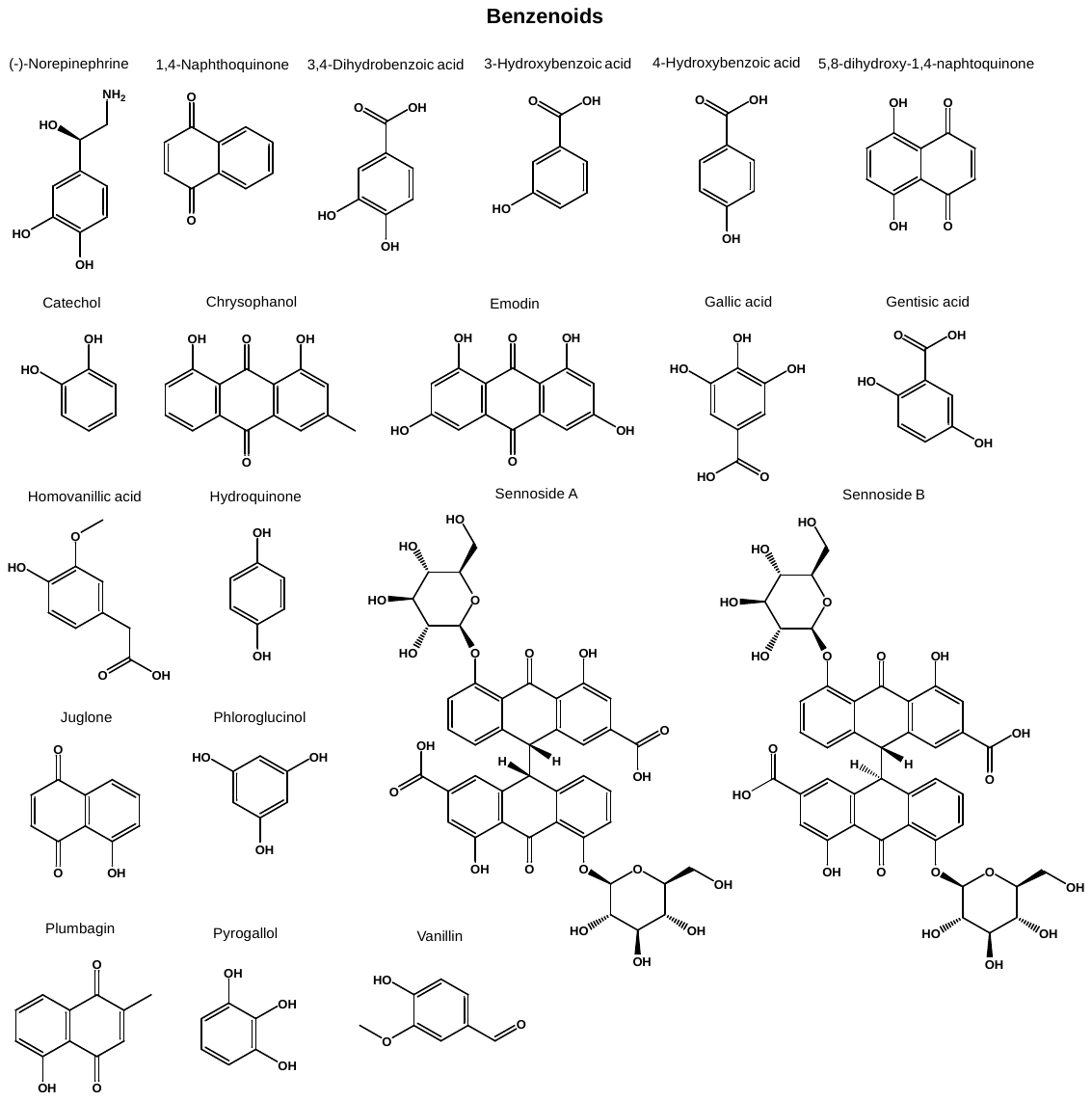

**Fig. S1 (cont.)**: Chemical structures of all tested molecules, grouped according to the respective chemical superclass according to Classyfire.

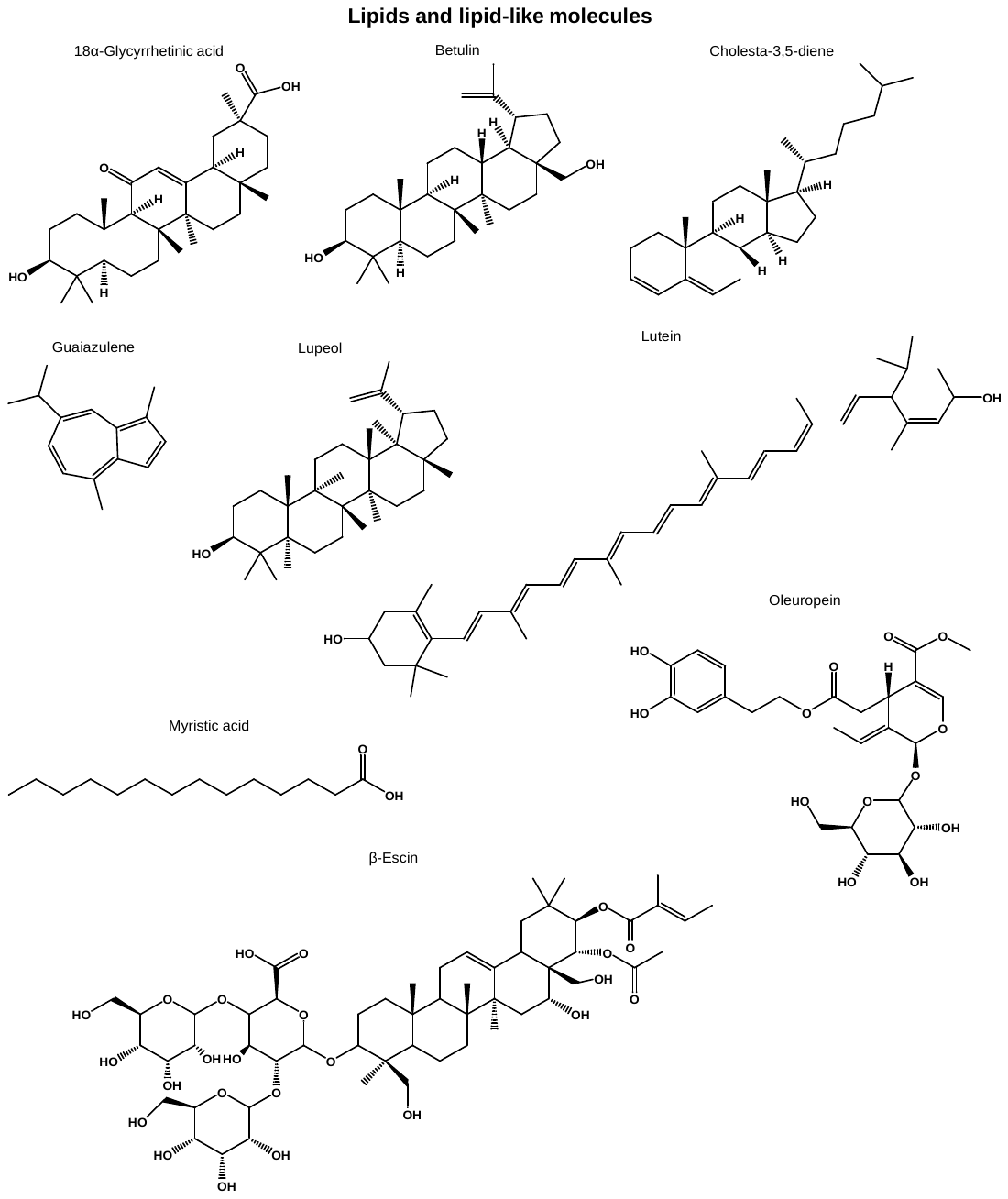

**Fig. S1 (cont.)**: Chemical structures of all tested molecules, grouped according to the respective chemical superclass according to Classyfire.

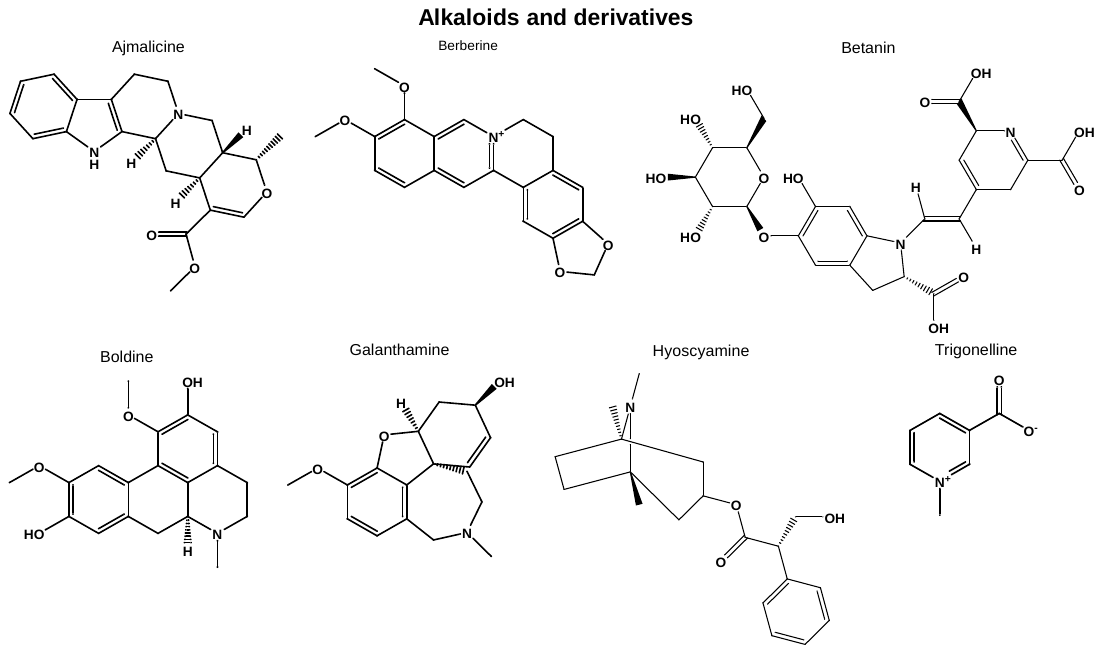

**Fig. S1 (cont.)**: Chemical structures of all tested molecules, grouped according to the respective chemical superclass according to Classyfire.

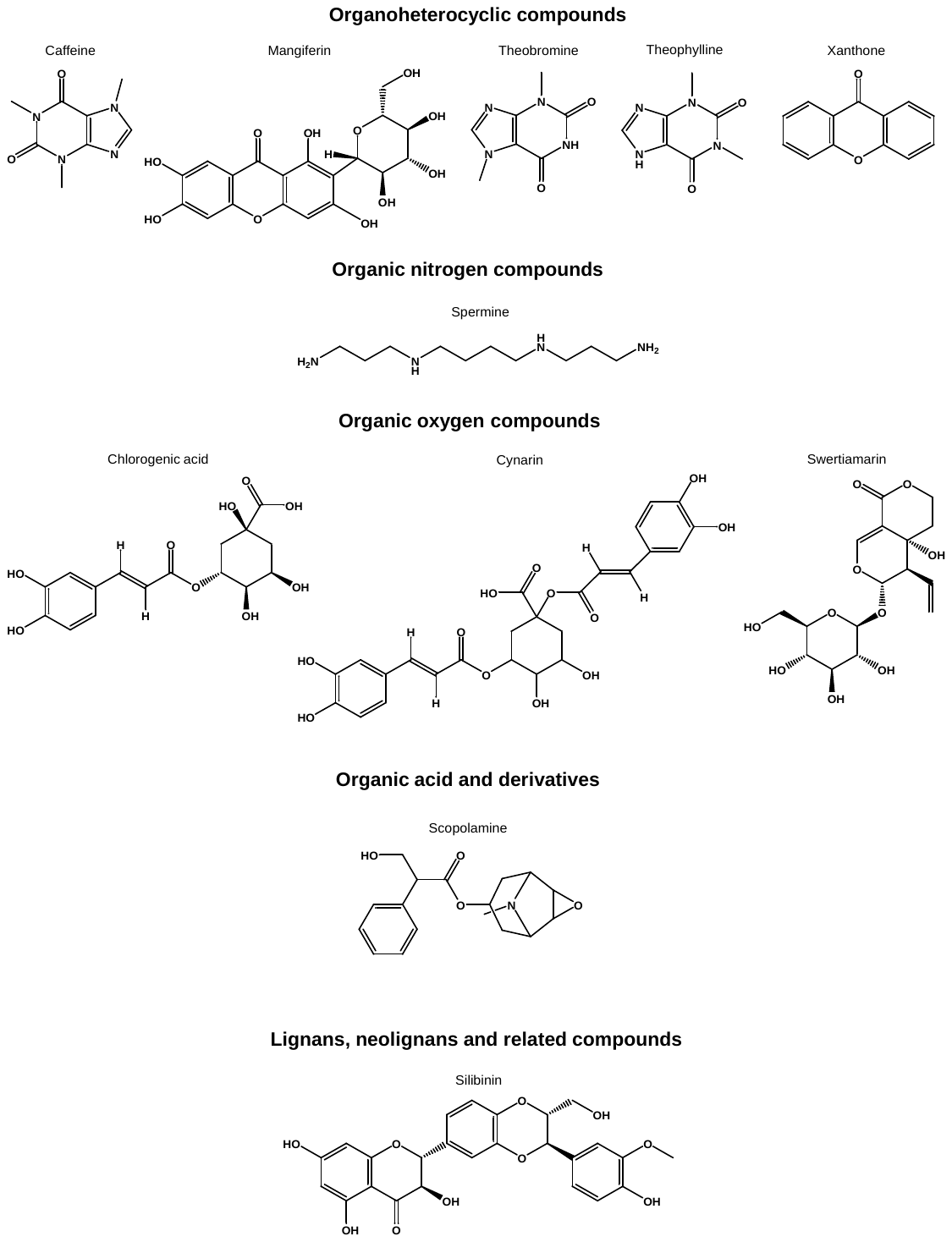

**Fig. S1 (cont.)**: Chemical structures of all tested molecules, grouped according to the respective chemical superclass according to Classyfire.

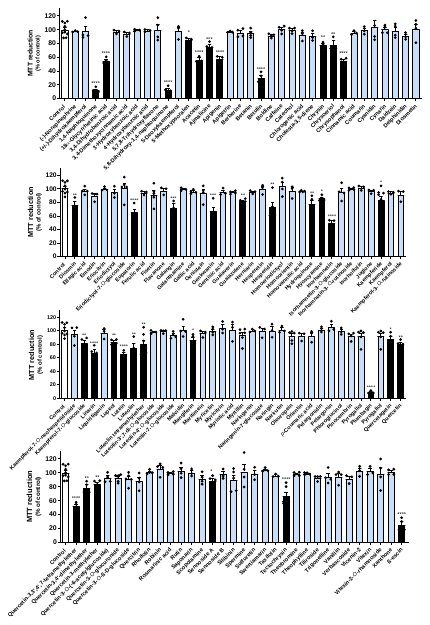

**Fig. S2.** Impact of a library of natural products on MRC-5 fibroblasts upon cell viability, as determined by MTT reduction assay. All compounds were tested at 50 µM. Results correspond to the mean ± standard error of the mean and represent, at least, three independent experiments, each performed in triplicate.

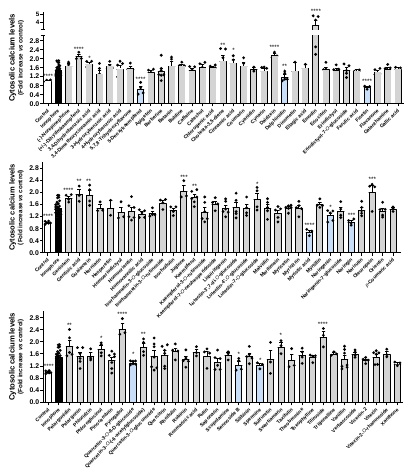

**Fig. S3.** Cytosolic calcium levels in MRC-5 fibroblasts in response to co-incubation of calcium ionophore A23187 at 5 µM and nontoxic molecules, as determined by the fluorescence of the probe Fura-2/AM. Results correspond to the mean ± standard error of the mean and represent, at least, three independent experiments, each performed in triplicate.

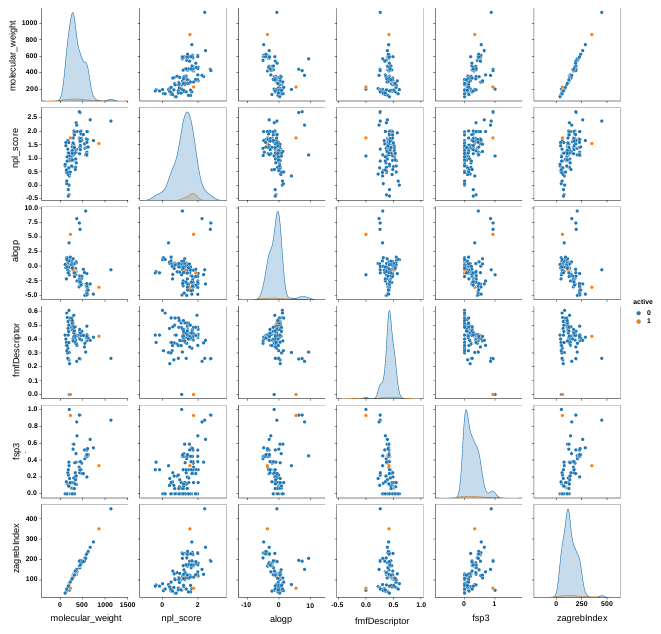

**Fig. S4.** Pairwise comparison of different features of the dataset. Blue = inactive molecules, orange = active molecules as *per* the results of **Fig. 5A**.

**Fig. S5.** Original images from which **Fig. 5D** and **6D** are derived. A: BiP expression on MRC-5 cells, B: GAPDH expression on MRC-5 cells, C: BiP expression on SH-SY5Y cells, D: GAPDH expression on SH-SY5Y cells.
